# Discovery of Z1362873773: A Novel Fascin Inhibitor from a Large Chemical Library for Colorectal Cancer

**DOI:** 10.1101/2024.08.13.606007

**Authors:** Alejandro Rodríguez-Martínez, Lucía Giraldo-Ruiz, Maria C. Ramos, Irene Luque, Diogo Ribeiro, Fátima Postigo-Corrales, Begoña Alburquerque-González, Silvia Montoro-García, Ana Belén Arroyo-Rodríguez, Pablo Conesa-Zamora, Ana María Hurtado, Ginés Luengo-Gil, Horacio Pérez-Sánchez

## Abstract

Metastasis is one of the leading causes of cancer-related death worldwide. Fascin is involved in this process by bundling actin filaments and producing protrusions in cancer cells, which facilitate their migration. It has been shown that the overexpression of this protein is related to the appearance of different types of cancer, such as colorectal cancer. In this study, we conducted an in silico screening against the enamine library, a compound library with a broad chemical space (≈1.4M compounds), followed by further validation with physicochemical assays and cellular migration and cytotoxicity tests, thereby obtaining a molecule with considerable fascin inhibitory and migration-arresting capacity similar to other inhibitors already known in the literature.

**Graphical Abstract:** 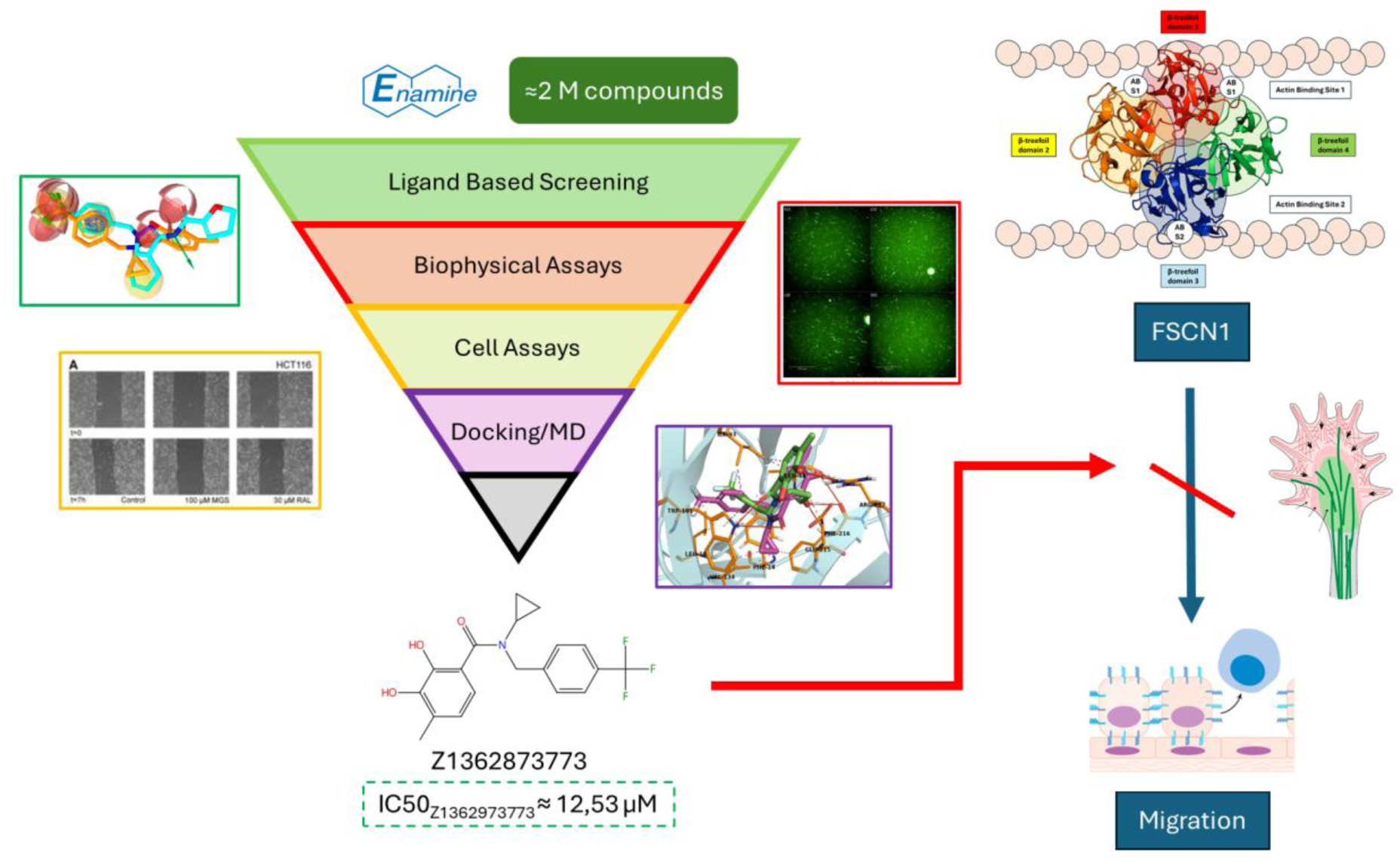

## Introduction

Cancer remains one of the most formidable challenges in the medical field, presenting a myriad of complex mechanisms that evade current therapeutic strategies and demand novel approaches to treatment^1^. Tumor metastasis is the primary cause of death from cancer^2^. Cancer cell migration and invasion are hallmarks of their metastatic capacity^3^, necessitating reorganization of the actin cytoskeleton. This reorganization facilitates the creation of protrusions, such as filopodia, lamellipodia, and invadopodia, which are instrumental in cancer cell movement^4^.

A pivotal protein in this process is fascin, which bundles actin filaments (F-actin) and is integral to developing protrusive structures. Unlike most normal epithelial tissues, where it is minimally expressed or not at all, fascin is notably increased in various human cancers. This elevation is associated with enhanced tumor growth, invasion, and metastasis^5,6^. Fascin^4,5^ has been identified as a promising therapeutic target and marker for aggressive cancer. Previous studies have shown its overexpression in different types of cancer, such as serrated adenocarcinoma (SAC), a colorectal carcinoma subtype recognized by the World Health Organization (WHO), due to its poor prognosis and more aggressive invasion, as evidenced by increased tumor budding, E-cadherin loss, and higher rates of KRAS or BRAF mutations compared to typical colorectal cancers^6–8^.

Migrastatin and its derivatives effectively inhibit fascin and have shown potential in reducing tumor cell movement, invasion, and subsequent metastasis^2^. However, the synthesis of these compounds is complicated by their complex structure, prompting the exploration of other anti-fascin agents, including those derived from indazol-furan-carboxamides^9^. Other molecules, such as the G2 compound and those derived from its chemical structure, have also demonstrated a strong response that inhibits fascin function^9^ .In addition, clear progress has been made in drug repurposing, obtaining some FDA-approved compounds, such as Imipramine and Raltegravir^10,11^.

However, FDA-approved compounds cover a minimal space concerning what could be studied, and there is still a chemical space that has yet to be explored in the search for novel fascin inhibitors. In addition, compounds obtained by repurposing are difficult to patent and obtain benefits in the short term. Thus, underexplored chemical compounds from large chemical synthesis libraries could be of great value and interest to biotech and pharmaceutical companies, highlighting the need for further exploration.

In this work, we meticulously conducted a computed-based screening of the Enamine HTS chemical library (1.368.754 molecules)^12^, followed by in vitro studies for the best potential hits. This comprehensive approach allowed us to assess the compounds’ effectiveness in halting cancer cell migration and invasion, providing a solid foundation for our findings.

## Materials & Methods

### Library generation

First, we selected the HTS compound library from Enamine to perform Ligand-Based Virtual Screening calculations. The library contains 1.368.049 compounds with a broad chemical space between them. The compound set was split into 217 files to allow parallelization and optimization of the corresponding calculations. From the command line, we launched the command *idbgen*(available under the LigandScout license) to generate an LDB format file for each section where the library was split.

### Pharmacophore model generation

The next step was generating a pharmacophore model for G2, an effective fascin inhibitor^10^. We obtained an SDF file with the inhibitor’s chemical structure from PubChem^13^ to obtain the model. The SDF file was uploaded to LigandScout GUI^14^, and the corresponding pharmacophore model was generated. Pharmacophore features that did not influence fascin binding or inhibition were removed. The model was saved in PMZ format to screen it against the Enamine library generated in the previous step.

### Ligand-based Virtual Screening by Ligand Scout

A massive Virtual Screening calculation was performed using the in-house software Metascreener, developed by our research group (https://github.com/bio-hpc/metascreener). The pharmacophore model was screened against all the generated Enamine libraries. The maximum number of features to be omitted (-a,-*allow_omit*) for this calculation was 0. The aim was to obtain all compounds whose pharmacophore features matched those of the G2 model. We applied a post-filtering step to retain only those molecules with a cutoff greater than 0.97 in pharmacophore similarity values. Finally, we selected the 12 available compounds in stock at Enamine from this subset to carry out in vitro assays.

### Biophysical characterization

The selected compounds were tested using two independent assays based on various principles. First, a primary assay was conducted using Differential Scanning Fluorimetry (Thermofluor), which assesses the compound’s ability to interact with the protein and induce changes in its denaturation temperature. Concurrently, validation was carried out using an F-actin-bundling assay with fascin.

### Differential scanning calorimetry (DSF) / thermofluor. Study of Binding to fascin

384-well plates were used for this assay, and a final assay volume of 10 μl was used. With the aid of Echo® Acoustic Liquid Handling (Beckman Coulter), 400 nL of each compound, from an initial stock of 10 mM and 100% DMSO, was dispensed into four wells to obtain four replicates of each at a final concentration of 400 μM. Two quality control measures were used for the assay: a control of native fascin, where the protein was present in DMSO at the same concentration as the compounds, and a control for inhibitory activity in the presence of BDP13176 at a concentration of 400 μM. Each control was placed in two complete plate columns, constituting 32 replicates for each condition. The thermal stability profile was measured using the CFX384 Touch Real-Time PCR Detection System (BioRad), and the Tm values in the native protein and the shift in the inhibition control and each compound were obtained. The ’Z-Factor’ was calculated as an informative parameter of the assay quality.

### F-actin bundling assay

We implemented an image-based assay to visualize F-actin bundling mediated by fascin cross-linking following the assay reported by Huang et al, 2014^15^. Labeling was performed with phalloidin conjugated to a commercial fluorescent probe (Alexa Fluor 488-Phalloidin, Thermofisher), directed at the positive charges of the F-actin filaments and bundles. Thus, F-actin structures can be visualized using fluorescence microscopy. The Operetta CLS High Content Analysis System (Revvity) was used to visualize and capture the images. Subsequently, the images were processed and analyzed using Harmony^TM^ (Revvity).

Each selected compound (300 nL) was dissolved in 100% DMSO from an initial stock (10 mM) using Echo® Acoustic Liquid Handling (Beckman Coulter). Next, an amount of 15 μL of pure fascin at a concentration of 0.5 μM in buffer (20 mM Hepes, 100 mM NaCl, pH 7.45) was dispensed onto a plate in a 384-well format (Corning) using an automatic dispenser (Multidrop Combi, Thermo Fisher), and allowed to incubate for 30 min. Subsequently, 15 μl of polymerized actin was added at a concentration of 0.5 μM (in 100 mM KCl, 20 mM Tris-HCl, pH 7.5, 2 mM MgCl2, 1 mM DTT, 1 mM ATP; Cytoskeleton Inc.). This resulted in a final concentration of 100 μM of the chemical compounds. After another 30 min of incubation, 10 μl of Alexa Fluor-488 Phalloidin (diluted 50-fold from 100% methanol stocks) was added to stain F-actin, and the mixture was incubated in the dark for 1 h. Next, 25 μl of the final solution was transferred to a 384-well plate coated with poly-d-lysine (Corning) and incubated for another 20 min before imaging.

Three quality control assays were performed. Two controls corresponded to negative controls for fascin cross-linking activity. These included F-actin in the absence of fascin to verify that actin bundling depends on the presence of fascin. Fascin at a 1:1 molar ratio was used as a positive control. BDP-1317615^16^, a known fascin inhibitor, was used at a concentration equal to that of the compounds tested. In each test, 16 repetitions were performed in different wells of each control.

### Cell culture

Two human colorectal adenocarcinoma cell lines (DLD-1 (CLS Cat# 300220/p23208_DLD-1, RRID: CVCL_0248) and HCT-116 (CLS Cat# 300195/p19841_HCT116.html, RRID: CVCL_0291)) were obtained from the American Type Culture Collection (ATCC, Rockville, Maryland, USA). The cell lines were cultured at 37 °C in high-glucose Dulbecco’s modified Eagle’s medium (DMEM) containing 10% heat-inactivated fetal bovine serum (FBS) and 50 μg/ml gentamicin (Cat#15750037 Thermo Fisher) in an atmosphere of 5% CO2 and 95% humidified air. Subcultures were performed when the cells reached 90% confluence. According to the American National Standards Institute, human cell line identification by short tandem repeat profile testing has shown an appropriate match between HCT-116 and DLD-1 cell lines.

### Cell viability assay

Exponentially growing cells were plated in quintuplicate in flat-bottomed 96-well plates (Nunc, Roskilde, Denmark) at 1500 cells/well and growth in a humidified 5% CO2 incubator at 37°C. The day after plating, drugs were serially diluted from 0.1 to 100 µM. Control wells contained medium without the drug plus 0.1% dimethyl sulfoxide (DMSO) (drug carrier). After 72 hours, cell viability was evaluated by adding 50 μl of activated XTT solution (Biotium, VWR, catalog number: 30007) to each well, and the plates were incubated at 37 °C for 4 hours^17^. Lastly, the absorbance at 450 nm was measured with a background subtraction reading at 630 nm, in a Spectramax ID3 plate reader (Molecular Devices, VWR).

### Organoids

Patient-derived colorectal cancer organoids were established and cultured in our Pathology Department using IntestiCult™ Organoid Growth Medium (Human) following the manufacturer’s protocol (Stemcell Technologies)^18^. The medium from each well intended for passage was removed without disturbing the domes; each well was then filled with 500 μl of ice-cold D-PBS, the solution was pipetted up and down to facilitate matrix breakage, and the suspensions were transferred to a 15 ml conical tube, which was repeated at least once to ensure complete retrieval of all organoids, with well checks conducted under an inverted microscope. Subsequently, the tubes were centrifuged at 290g for 5 min at a temperature between 2-8 °C, the supernatant was discarded, without disturbing the organoid pellet, and then treated with 1 ml of TrypLE™ Express Solution and incubated for 5 min at 37 C. After incubation, 100 μl of FBS was mixed with the suspension, the tubes were centrifuged at 250 g for 5 minutes at 2-8 °C, and the supernatant was discarded. To the obtained pellet, 1000 μl of DMEM/F-12 + 1% BSA was added and the number of organoids in the suspension was counted using a Countess™ 3 Automated Cell Counter with Countess Reusable Slides (Invitrogen, Thermo Fisher Scientific). Considering that the domes of the 96-well plate corresponded to 5 μl and each dome had 5000 cells, the correct amount of suspension to be used for the assay was calculated, always considering the 1:1 ratio of cell suspension and Matrigel® and that each treatment/condition was performed in quadruplicate. Pre-wetting the pipette tips with cold DMEM/F-12 + 1% BSA before manipulating the cell suspension was used to prevent sticking to the wall of the pipette tip, and to place the 96-well plate in the incubator for at least 30 min before starting the assay. After plating, the plates were placed in an incubator at 37 °C and 5% CO2 for 15 min to allow the solidification of the domes. Then, approximately 100 μl of D-PBS was added to the wells in the borders to avoid evaporation of the cell culture medium, and 50 µL of complete IntestiCult™ Organoid Growth Medium at room temperature (without Y-27632 and gentamicin) was added. After 48 h, the control with the drug carrier (0.1% DMSO) and the respective drugs in serial dilutions from 0.1 to 100 µM were added to the corresponding wells (in quadruplicate) at a total volume of 100 μl per well. Subsequently, the plates were left in the incubator for 5 days, and photos of the wells were taken every day using an inverted microscope, as a way to register the growth and the treatment effect on the organoids. On the fifth day, cell viability was determined using the CellTiter-Glo® Luminescent Cell Viability Assay (Promega) following the manufacturer’s guidelines. Luminescence was measured using a Spectramax ID3 plate reader (Molecular Devices, VWR)^19–23^.

### Molecular modeling

We conducted molecular modeling of fascin structure to determine the interaction between the predicted compounds and fascin. First, we used PDB code 6B0T, which corresponds to the binding between the crystallographic structure of fascin and NP-G2-029, a derivative of G2.

The structure was preprocessed using MAESTRO’s Schrödinger software^24^. Chain A was extracted, and the corresponding hydrogens and charges were added to the structure. The new structure was saved in the mol2 format. A similar protocol was used with AutoDockTools^25^ software, which performed the same tasks and applied the AD type to the atoms. In this case, the structure was saved in the pdbqt format.

The predicted ligands were converted to mol2 and pdbqt formats using ChemAxon (MolConvert tools)^26^ and AutoDockTools, respectively. In each case, corresponding charges were added.

### Blind Docking

When the protein and candidate ligand files were processed, two Blind Docking calculations were performed using the NP-G2-029 Fascin (PDB:6B0T)19 and free protein (PDB:3P53)20 structures to explore the conformational space and the corresponding possible binding site of the predicted compound. For each Blind Docking, we used two docking software packages: Lead Finder^27^ and AutoDock Vina^28^. A consensus between the two methods was achieved to determine the average pose.

### Targeted Docking

Targeted docking was performed in the actin-binding site 2 coordinates, where NP-G2-044 binds to fascin in the 6B0T PDB structure^29^. The pose of the best clusters was determined. Lead Finder and AutoDock Vina were used, with the respective consensus between both. In the next step, we chose the pose obtained by the Lead Finder for the molecular dynamics (MD) simulation.

### MD simulations

To validate the stability of the complex in an aqueous system and whether the binding to actin-binding site 2 was maintained over time, we performed an MD simulation of the fascin-Z1362873773 complex in the binding area with the pose obtained by the Lead Finder virtual screening calculations. We also ran an MD simulation for the crystallized fascin in complex with NP-G2-029 (6B0T) to compare the evolution of Z1362873773 with NP-G2-029, which is a ligand whose binding to actin-binding site 2 has been more extensively studied.

We performed Molecular Dynamics simulations at 100 ns on the same fascin structure 6b0t for the poses obtained by the BD calculations, corresponding to the known binding site actin-binding site 2^29^, with the best docking score in the blind docking. In addition, we performed an MD simulation for the NP-G2-029 and fascin complex using the crystallographic structure (PDB: 6B0T) to compare the results with a complex with a ligand bound in the same site^29^. For this, we first generated the topology of the ligand for each pose using an automatic script that utilizes ACPYPE^30,31^

We followed the subsequent steps of the molecular dynamics simulations using GROMACS 2022.3^32^. These simulations were launched on the Picasso server (https://www.scbi.uma.es/web/es/inicio/) using a GPU (NVIDIA A100-SXM4-40GB) and 4 GB of RAM. We first created the protein topology using the gmx pdb2gmx command, specifying AMBER99SB as the force field. The simulation box was defined as the solvated and added ions. The next step was an energy minimization stage of 2000 ps. After that, a single NvT equilibration stage of 50000 ps and five NpT equilibration stages of 50000 ps each were carried out. Finally, the dynamics were run at 100 ns, and the final trajectory generated was extracted in different frames to compare ligand stability and movement concerning the binding pose.

### Trajectories analysis

The results obtained by MD simulation were analyzed using ASGARD (https://github.com/bio-hpc/ASGARD), an in-house tool developed by our group^33^. We calculated the non-bonded interactions and hydrogen bonds between the final ligands. In addition, we studied the stability of the protein-ligand complex and the flexibility and dynamics of the fascin produced by its interaction with the binding site. The required input files were prepared for the tool, and plots and raw data were generated.

### Drug-like property predictions

Drug-like properties were calculated using the MAESTRO Schrödinger tool QikProp^34^. The properties were obtained for the most known fascin inhibitors, imipramine, migrastatin, raltegravir, NP-G2-044, G2, BDP-13176, and the Z1362873773 predicted compound. The values were saved in a CSV file. A comparative table was generated with this data. A Ligand-Based screening for two models created from the Z1362873773 compound against the DrugBank compounds database was performed to study possible alternative targets where the hit molecule could act. The first model was created by calculating all the possible features of the molecule, whereas the second model only had the Z1362873773 features that matched the G2 pharmacophore model. For these calculations, the maximum number of features omitted was four for the first model and one for the second.

## Results

### Ligand-based in silico screening

We obtained 212 compounds with all features matching the pharmacophore model for G2 generated by LigandScout (Figure 1). Those molecules with a Relative-Pharmacophore score higher than 0.97 were chosen for subsequent steps. Finally, the 12 compounds selected (Z1362873773, Z335377818, Z29860525, Z335377814, Z735522228, Z62748384, Z107323688, Z44912418, Z1229770825, Z79431411, Z1797081400 and Z237374404) was tested in vitro in the study’s next step. Figures 2 and 3 show a graphical comparison of the pharmacophore model for G2 with the Z1362873773 Enamine compound, the one with the best in vitro results.

**Figure 1.**
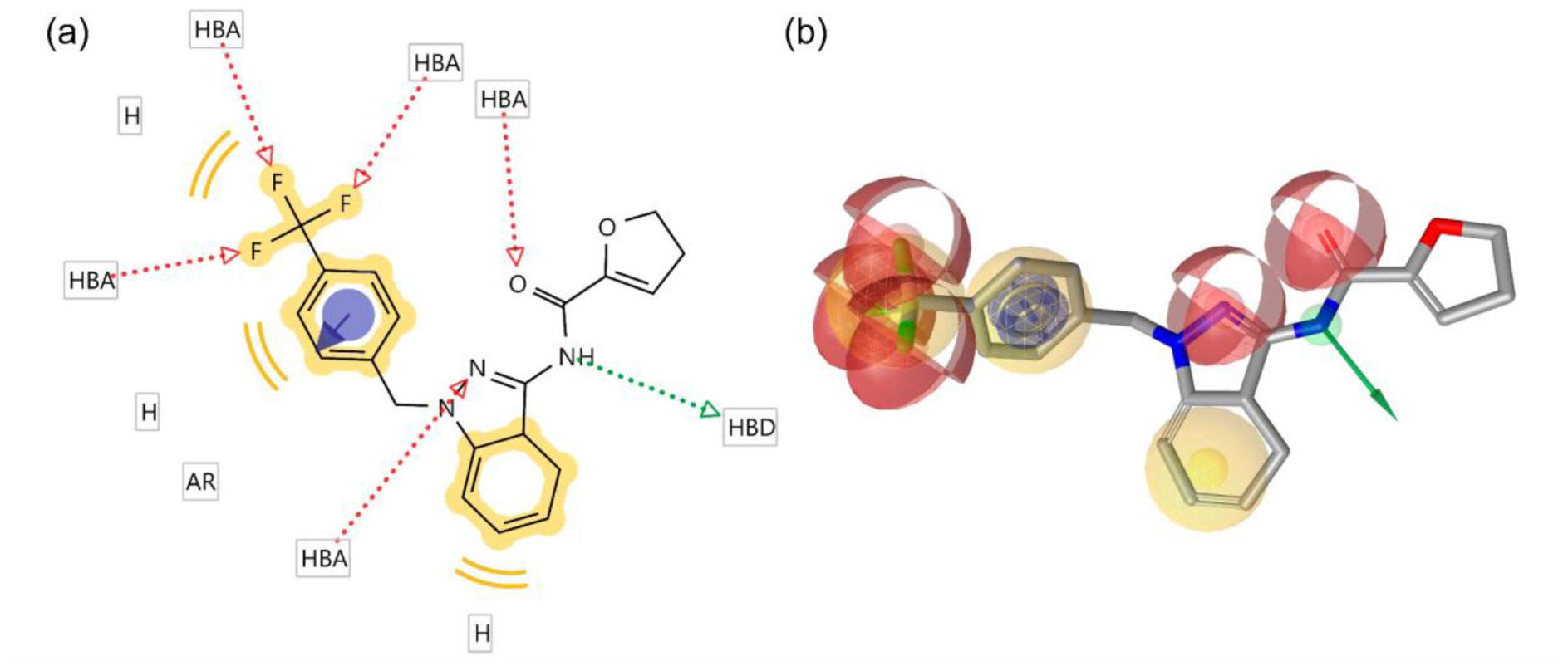
(a) 2D representation of the pharmacophore model for the G2 compound generated by LigandScout (b) 3D representation of the pharmacophore model for the G2 compound generated by LigandScout. HBA (red): hydrogen bond acceptor; HBD (green): hydrogen bond donor; H (yellow): hydrophobic.

**Figure 2.**
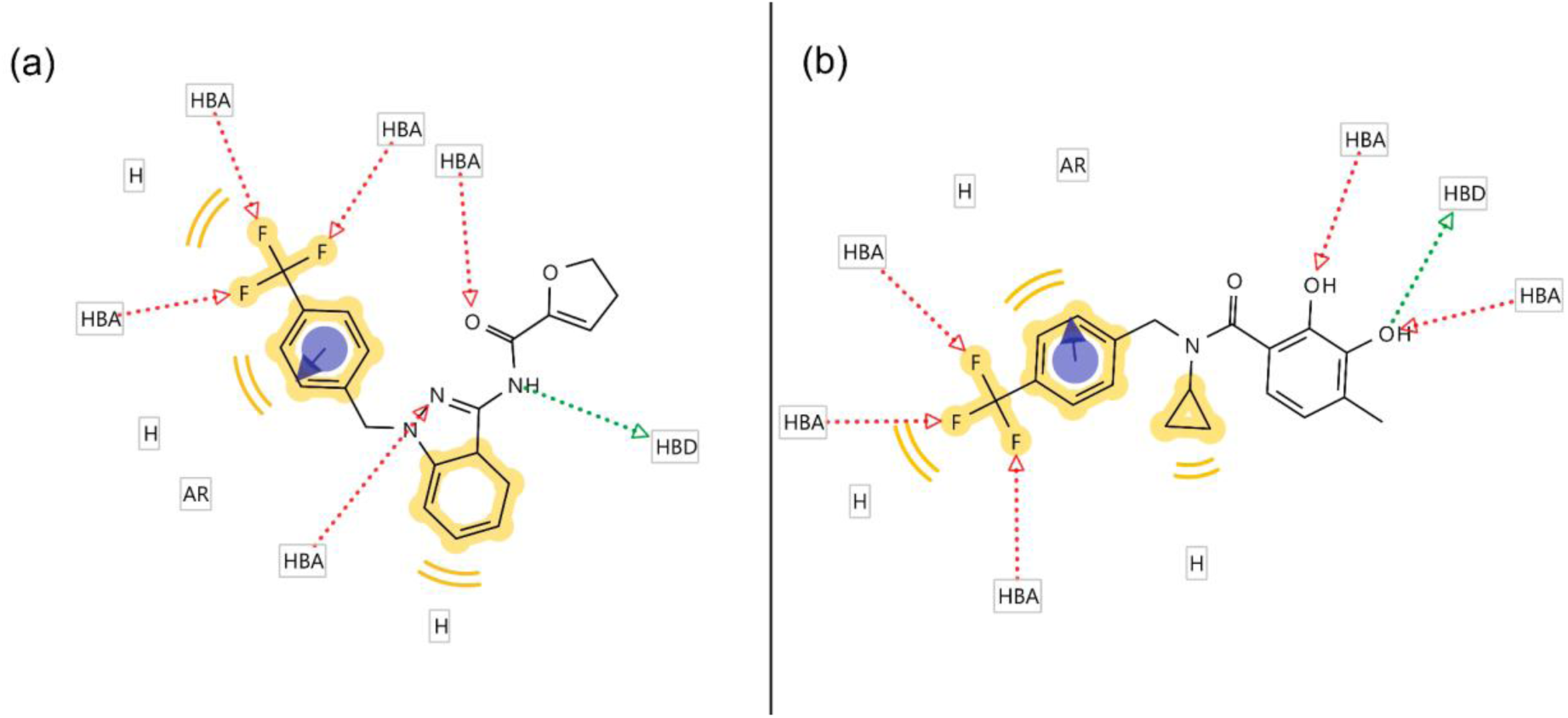
Comparison between the chemical structure and pharmacophore model of the G2 compound (a) and the compound obtained via VS from the Enamime library (Z1362873773).

**Figure 3.**
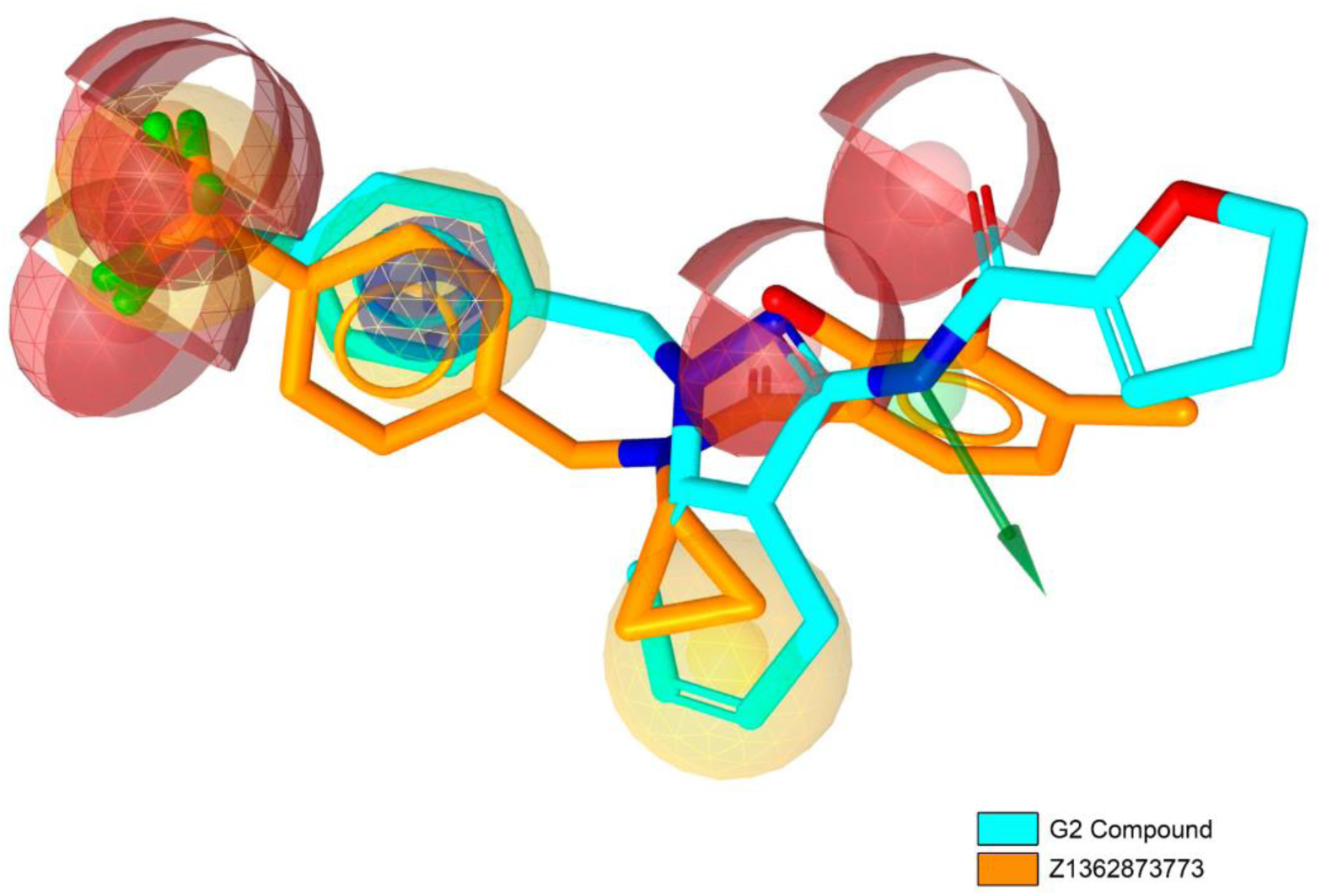
Comparison between the tridimensional chemical structures of the G2 compound (cyan) and the Enamine compound obtained by Virtual Screening, Z1362873773 (orange).

**Figure 4.**
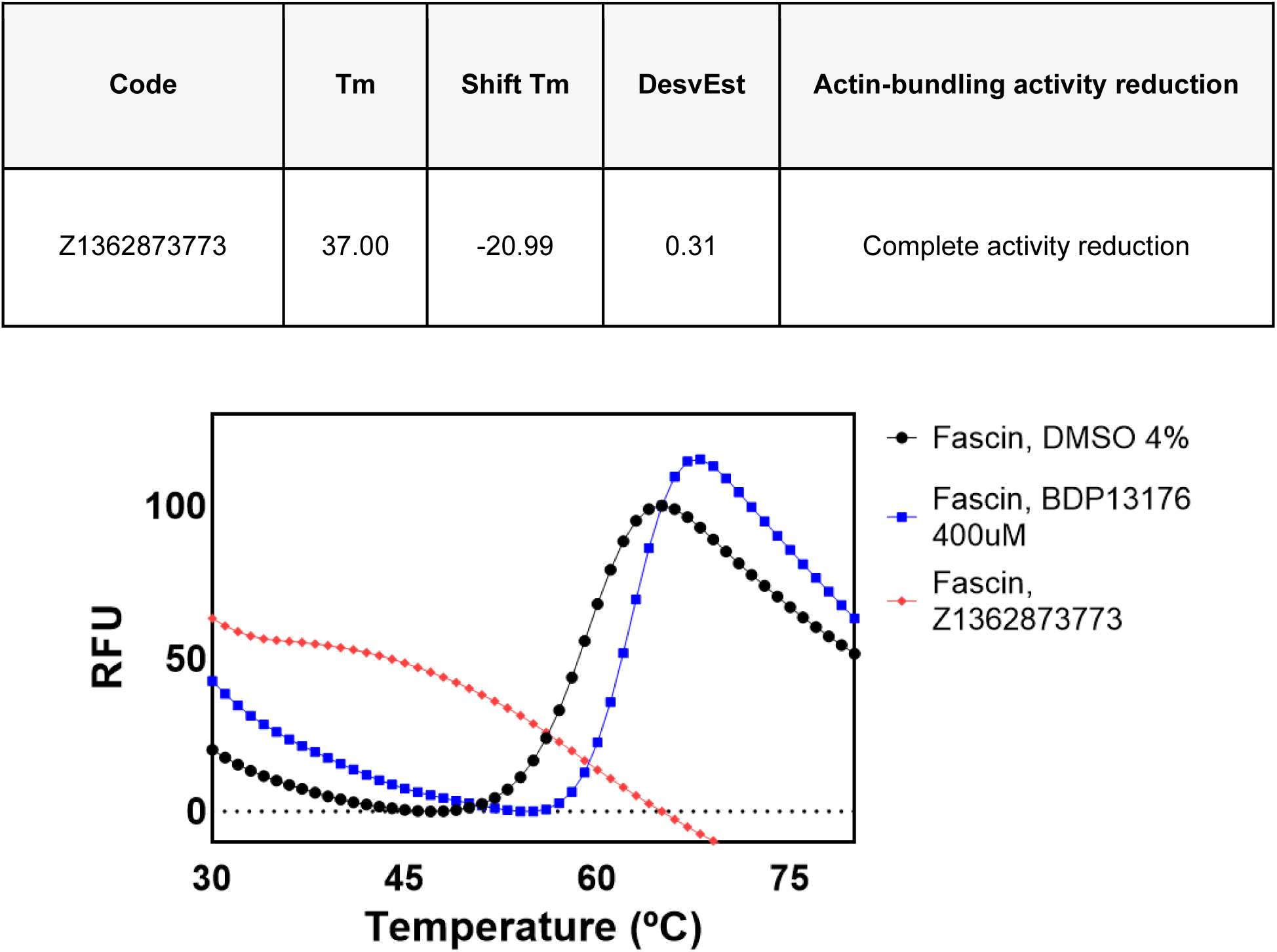
Denaturation curves of wild-type fascin in the absence of an inhibitor (red), in the presence of BDP-13176 (blue), and in the presence of the pure compounds tested (highlighted in black).

### Thermofluor assays validate Z1362873773 binding to fascin in vitro

The ability of the selected compounds to bind fascin was evaluated using differential scanning fluorimetry. At pH 7.4, fascin unfolds through a singular transition without showing any dependency on concentration and exhibits a high level of DMSO tolerance. The average melting temperatures (Tm) for unbound fascin were recorded using different internal filters (FAM, HEX, and T-Red), serving as a reference to assess Tm changes upon compound binding (Tm,FAM = 55.6 ± 0.5 °C, Tm,HEX = 56.0 ± 0.0 °C, Tm,Tred = 56.2 ± 0.6 °C, with error values representing the standard deviation for the seven replicates used as internal controls on each plate). Some compounds led to altered thermal unfolding profiles, making it difficult to accurately determine Tm values, likely due to the compounds interfering with the SYPRO fluorescence signal. Among those analyzed, Z1362873773 showed the most significant shift, reducing Tm 20.99°C, indicating its specific binding to fascin. This considerable deviation suggests a potential destabilizing effect on the protein structure induced by the compound.

### Z1362873773 reduces the fascin bundling activity

Z1362873773 compound was assayed for the fascin bundling activity by imaging-based assay. F-actin bundles bear negative charges that are captured by a positively charged amino acid polymer, poly-D-Lysine, which coats the well surface of the plates, then fascin cross-links the F-actin filaments into straight, compact and rigid bundles. When an inhibitor of fascin is present in the well, the F-actin filaments are unstructured. By labelling F-actin with phalloidin-conjugated with fluorescent dye, we could visualize the F-actin structures using a high-content imaging system. Z1362873773 compound was assayed at 400 µM and total inhibition of F-actin bundles was observed (Figure 5), confirming the results obtained in Thermofluor assay.

**Figure 5.**
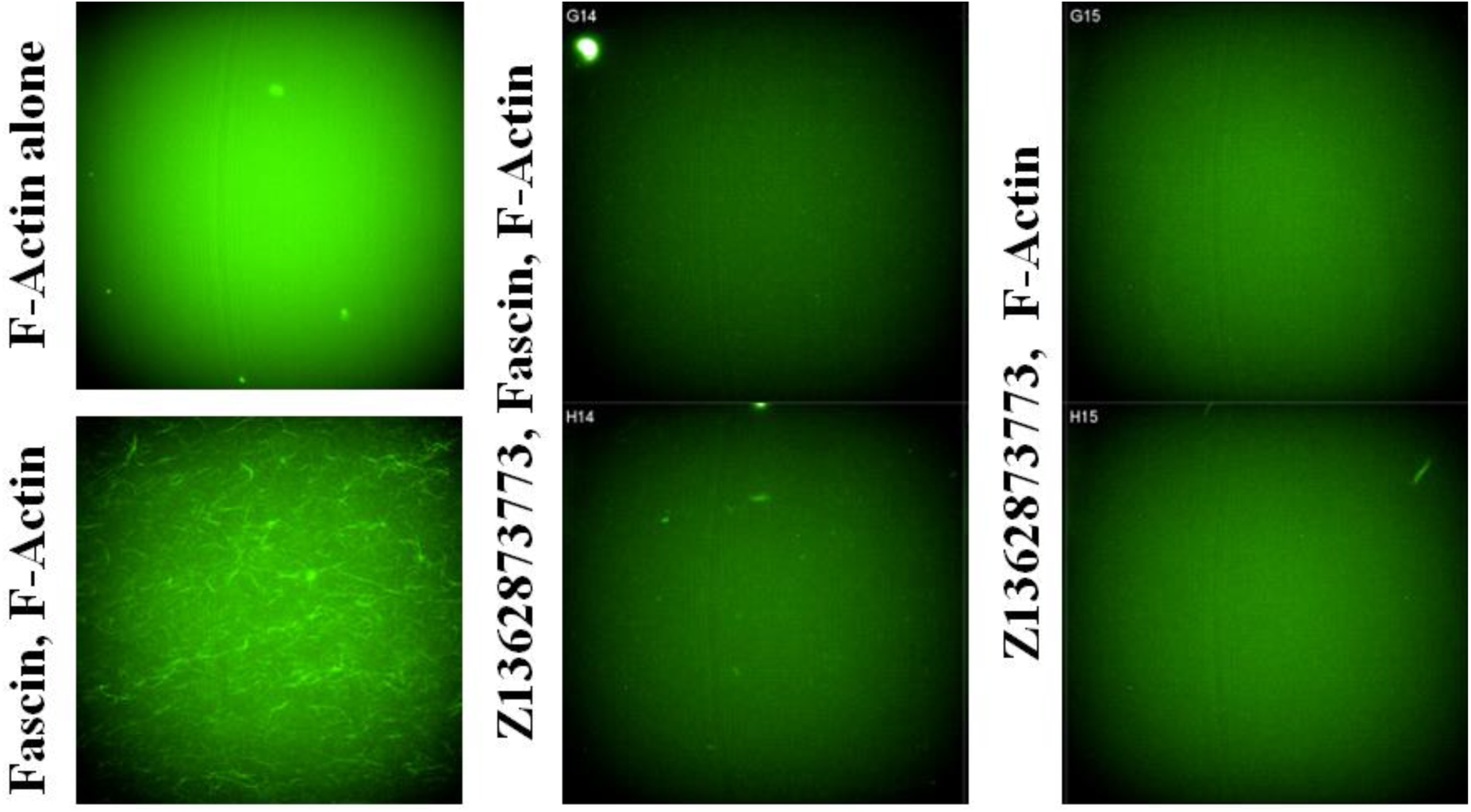
The formation of F-actin filaments was inhibited in the presence of the 400 µM Z1362873773 compound. The images showed are those of controls, F-actin plus fascin and F-actin alone, 400 µM BDP-1317615 (positive control of inhibition), and Z1362873773, with and without fascin, indicating that the bundling activity of fascin is likely to be specifically inhibited.

**Figure 6.**
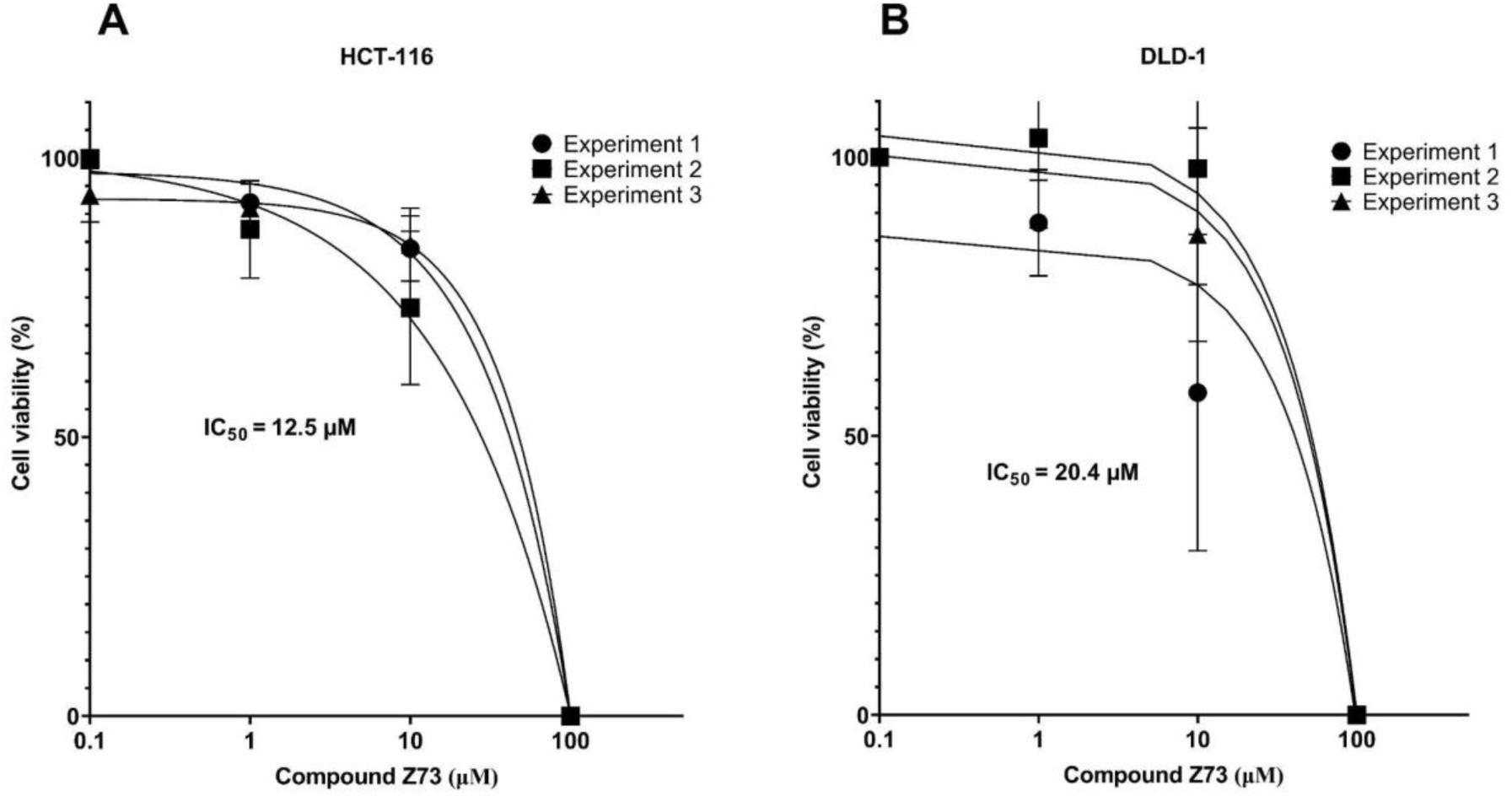
Cell viability assays for HCT-116 and DLD-1 cells were performed in three independent experiments. Each line represents the normalized curve fit for an independent experiment. Each point was determined in quintuplicate (mean value + SEM).

### Z1362873773 affects the cancer cell viability

Viability assays were performed on DLD-1 and HCT-116 cells to determine the drugs’ working concentrations. HCT-116 was more sensitive than DLD-1, and the working concentrations were set for subsequent *in vitro* studies at 12.5 and 20.4 µM Z1362873773.

### Z1362873773 inhibits the cancer cells’ migration

To investigate the impact of Z1362873773 on cell migration, cells treated with Z1362873773 were analyzed for motility using a wound healing assay at 24 and 48h. As shown in Figure 7, 20 µM Z1362873773 significantly reduced the migration of all tested cell lines (p < 0.05). The inhibitory effect of Z1362873773 was similar to that of G2, at the same concentration. Figure 7 a and b show the detailed decrease in the migration of HCT-116 cells at 24 and 48 h, respectively. Figure 7 c and d shows a detailed decrease in the migration of DLD-1 at 24 and 48h, respectively.

**Figure 7.**
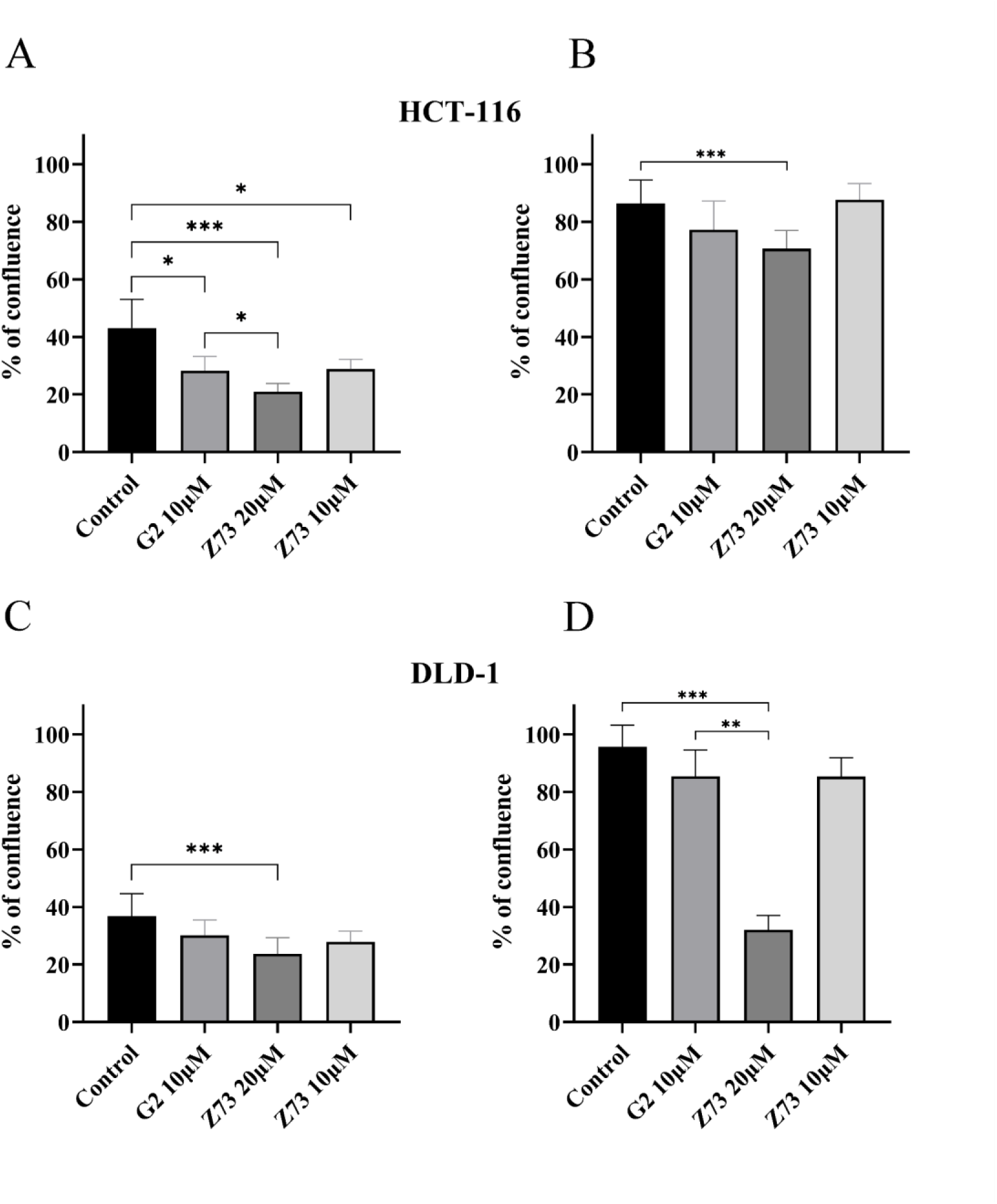
Wound healing assays for HCT-116 (A and B) and DLD-1 (C and D) cells at 24 and 48h using G2 and Z1362873773.

### Organoids

Viability curves in the presence of G2 or Z73 for primary (107T and 114T) and metastatic (06M) colorectal cancer organoids. Compound Z73 showed a 2-3 times significantly higher cytotoxic effect in our ex vivo model compared to G2 (Figure 8).

**Figure 8.**
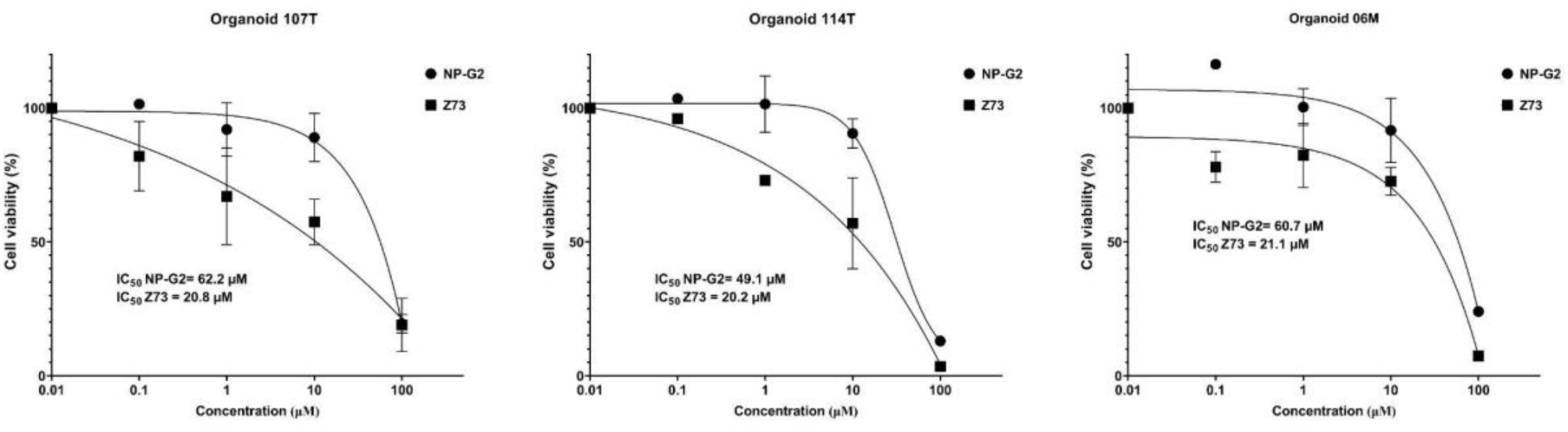
Viability curves in the presence of G2 or Z73 for primary (107T and 114T) and metastatic (06M) colorectal cancer organoids.

### Blind Docking finds a top pose in the actin-binding site 2

We performed two docking experiments using two different fascin structures. 6B0T corresponds to the structure obtained when NP-G2-029 (G2 analog) binds to actin-binding site 2 (Figure 9A). Blind Docking using AutoDock showed the pose inside the NP-G2-029 binding site as the first cluster with -8.9 kcal/mol. In the LeadFinder method, Blind Docking calculations were also performed in the free protein structure (PDB:3p53) to explore the structural flexibility of fascin. In the AutoDock Vina results, the poses for the top three and four clusters were found close to the position of actin-binding site 2, and the third cluster pose interacted with Phe216, one of the key residues for actin-binding site 2. In the results obtained from LeadFinder (Figure 9B), the first and second clusters were located at actin-binding site 2, exhibiting binding energies of 8.54 and -7.39 kcal/mol, respectively. Notably, cluster 1 also interacted with Phe216 through a hydrogen bond.

**Figure 9.**
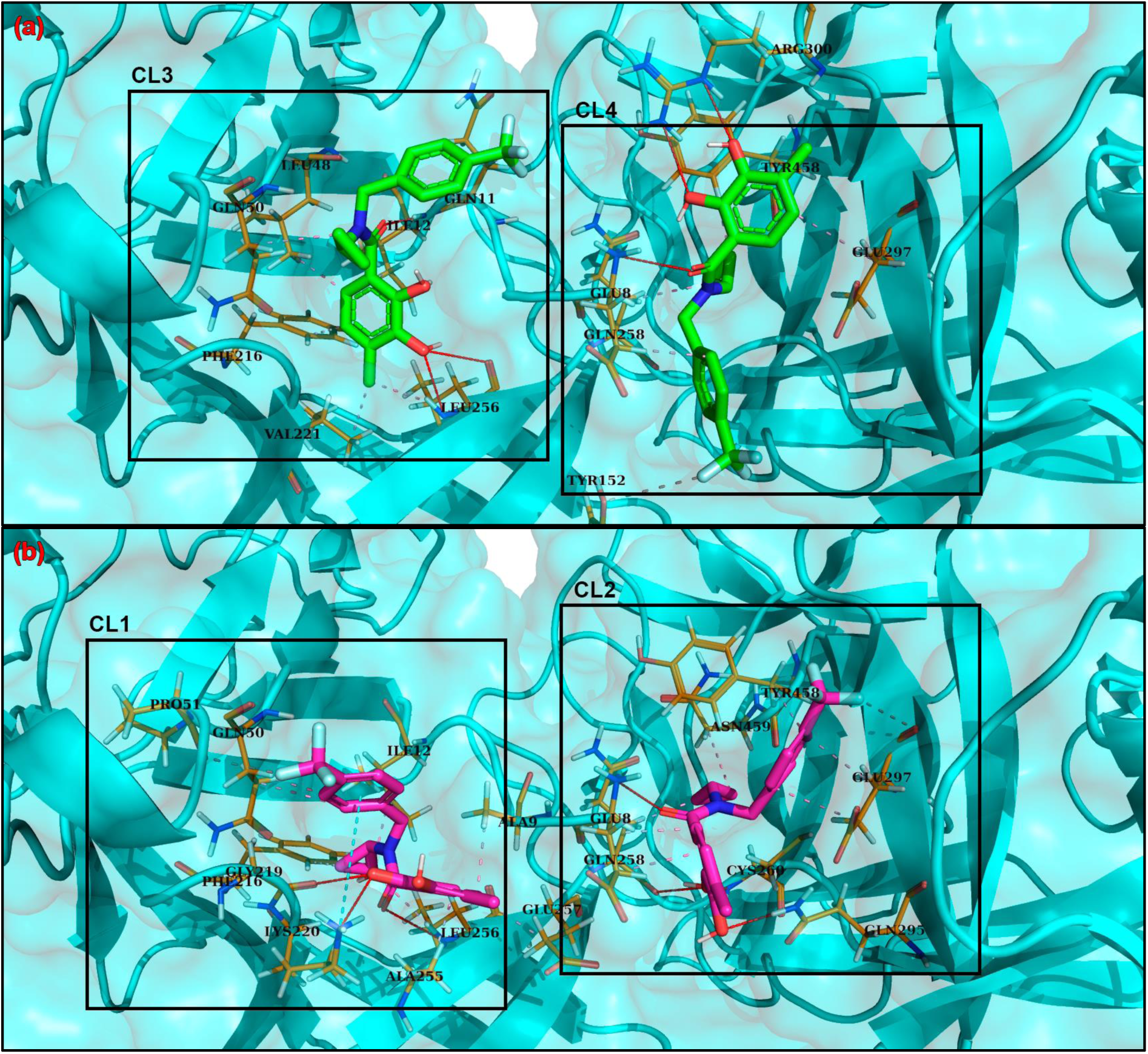
Poses obtained by BD calculations for fascin-free proteins (PDB:3P53). (a) Poses obtained using AutoDock Vina. The poses obtained represented the top three and four clusters, respectively. (b) Poses obtained using LeadFinder. The poses obtained represent the top one and two clusters. Regarding the Blind Docking calculations performed for the 6b0t structure, using AutoDock Vina (Figure 10A) and LeadFinder (Figure 10B) methods, actin-binding site 2 was detected in the top 1 cluster. The binding energy values were 8.92 and 8.72 kcal/mol for AutoDock Vina and Lead Finder, respectively. Furthermore, the key residues in the binding site were identified in both cases. We found hydrogen bond interactions between the compound and Phe216 and other hydrophobic interactions with crucial residues such as Ile93, Trp101, and Phe14^29^.

**Figure 10.**
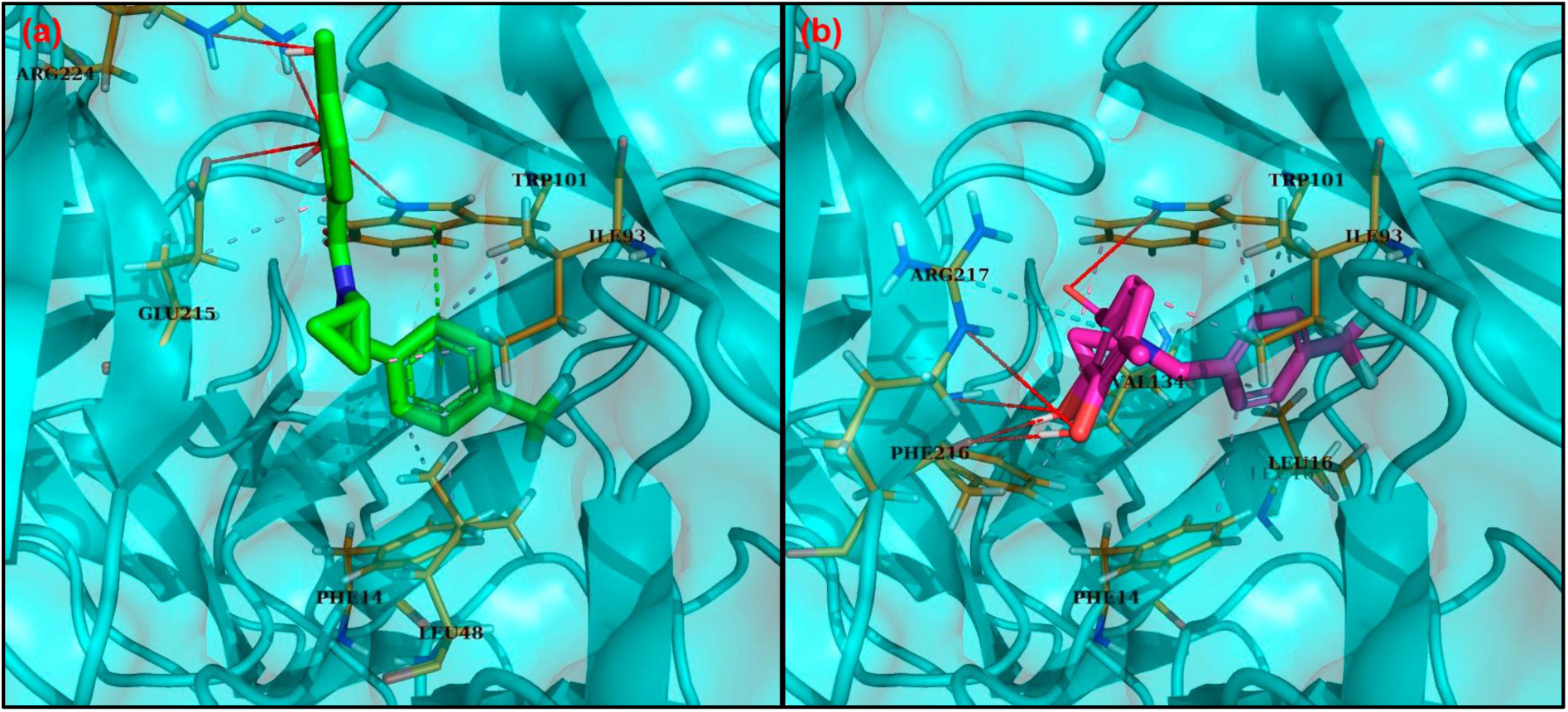
Poses obtained by BD calculations for fascin-NP-G2-029 (PDB ID:6B0T). (a) Pose obtained using AutoDock Vina. The obtained poses represented the top 1 cluster. (b) Pose obtained by LeadFinder. The poses obtained represented the top 1 cluster.

### Target Docking shows several interactions with actin-binding site 2 key residues

The Blind Docking calculation demonstrated that actin-binding site 2 is a potential binding site for the compound. Next, we carried out target docking in that position in the 6B0T structure (corresponding to the NP-G2-029 fascin complex). The AutoDock Vina and LeadFinder techniques obtained docking scores of -8.35 and -8.27 kcal/mol, respectively. Subsequently, we determined the specific interactions found using each method (Table 1). The interactions between the compound and the specific fascin residues in the binding site 1 found by AutoDock Vina calculations were the following: Phe14 (3.53 Å), Leu48 (3.80 Å), Ile93 (3.66 Å, 3.30 Å) and Glu215 (3.48 Å) for hydrophobic interactions; Trp101 (2.81 Å) and Arg217 (3.94 Å) for hydrogen bonds; and Trp101 (4.76 Å) for π-Stacking. Regarding the interactions obtained by LeadFinder software, we found hydrophobic interactions between the ligand and the residues Phe14 (3.82 Å), Leu16 (3.37 Å), Ile93 (3.33 Å, 3.27 Å), Trp101 (3.15 Å, 3.39 Å), Val134 (3.67 Å, 3.58 Å) y Phe216 (3.06 Å). LeadFinder also detects hydrogen bonds interaction with Trp101 (4.06 Å), Phe216 (3.00 Å, 3.04 Å, 2.81 Å) y Arg217 (3.41 Å).

**Table 1.** Interactions between Z1362873773 and fascin residues were detected by AutoDock Vina and Lead Finder calculations.

| Software | Docking Score (kcal/mol) | Hydrophobic Interactions (Å) | Hydrogen Bonds (Å) | π-Stacking Interactions (Å) | π-Cation Interactions (Å) |
| --- | --- | --- | --- | --- | --- |
| AutoDock Vina | -8.35 | **Phe14** (3.53),<br>**Leu48** (3.80),<br>**Ile93** (3.66, 3.30),<br>Glu215 (3.48) | **Trp101** (2.81),<br>Arg217 (3.94) | **Trp101** (4.76) | Arg217 (4.86) |
| LeadFinder | -8.27 | **Phe14** (3.82),<br>**Leu16** (3.37),<br>Ile93 (3.33, 3.27),<br>**Trp101** (3.15, 3.39),<br>**Val134** (3.67, 3.58),<br>**Phe216** (3.06) | **Trp101** (4.06),<br>**Phe216** (3.00, 3.04, 2.81),<br>Arg217 (3.41) | - | - |

Figure 11, a PyMOL session, visually represents the results and displays both poses obtained by AutoDock Vina and Lead Finder in the same fascin structure, allowing for a clear comparison. The residues obtained in each case matched some of the key residues at the actin-binding site 2^29^, further confirming the accuracy of our findings.

**Figure 11:**
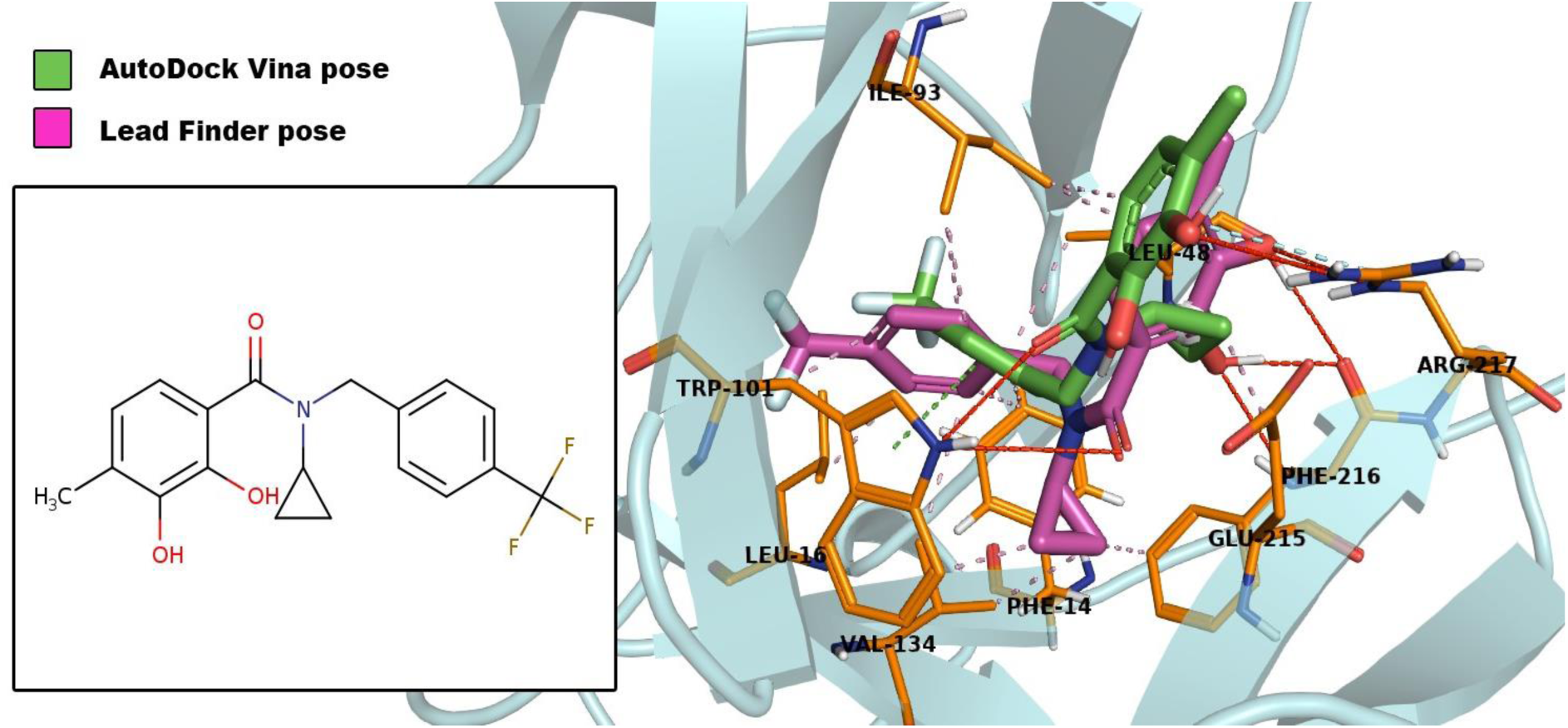
Pymol session showing the interactions between compound Z1362873773 and the crystallographic structure for fascin (PDB:6B0T) obtained by the docking calculation using AutoDock Vina (green) and LeadFinder (purple). In bold, these residues match the known critical residues in the interactions with the fascin structure. Pink interactions show hydrophobic interactions, red-colored interactions indicate hydrogen bonds, green-colored interactions are π-stacking, and cyan-blue-colored interactions correspond to π-cation interactions.

### MD analysis shows the Z1362873773 stability in the actin-binding site 2 pocket

We performed an MD simulation of Z1362873773 to study the stability of the ligand in the complex over time and in an aqueous system, considering fascin flexibility. In addition, we ran another MD simulation for the NP-G2-029 Fascin complex as a comparative test with a known fascin inhibitor bound to the actin-binding site 2 system.

The initial focus was on the RMSD during the simulation for the protein as the ligand, a key indicator of the system’s stability in both cases. Figure 12 illustrates the stabilization of the ligands in each complex, with both NP-G2-029 and Z1362873773 RMSD values maintaining a consistent range of 0.1 to 0.2 throughout the MD. The fascin structure, a known flexible protein, demonstrated significant flexibility during the simulation time, as expected.

**Figure 12:**
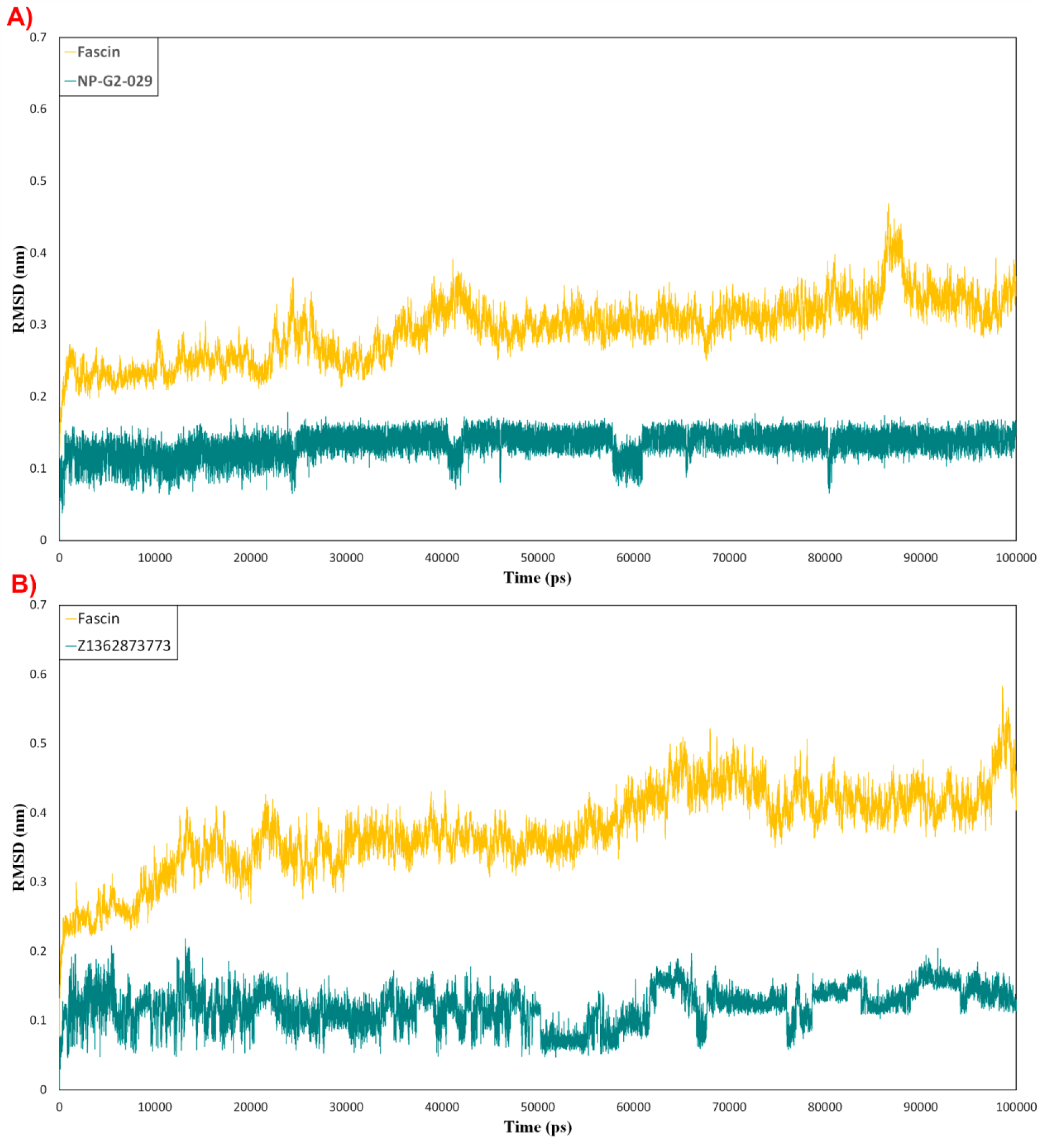
RMSD for protein (amber color) and ligand (teal color) in the NP-G2-029-fascin (A) and Z1362873773-fascin (B) complexes during the MD simulation. Molecular Mechanics Poisson-Boltzmann Surface Area (MMPBSA) analysis estimates the binding free energy between a protein and its ligand. We performed MMPBSA calculations for both complexes to examine the stability and strength of the binding between NP-G2-029 and Z1362873773 in actin-binding site 1 of fascin. The NP-G2-029 fascin complex stabilized the binding energy from the first 20 ns to between -250 and -300 kJ/mol (Figure 13A). Regarding the Z1362873773 compound, the binding energy of its complex with fascin started to stabilize in the first 10 ns, remaining at binding energy values of -200 and -250 -kJ/mol2 (Figure 13B). These analyses also demonstrate that the van der Waals contributions are more prominent than electrostatic interactions.

**Figure 13.**
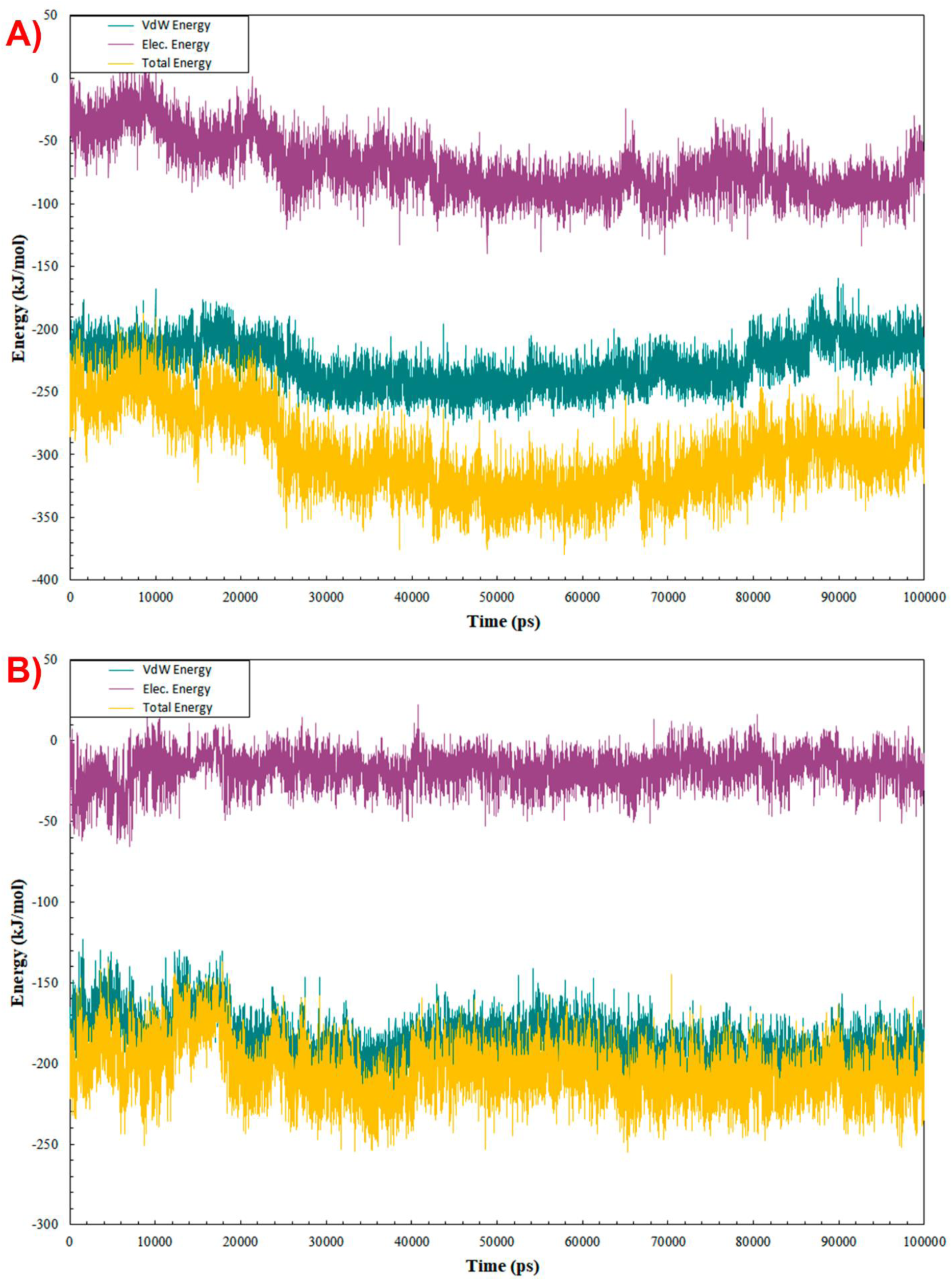
MMPBSA analysis of NP-G2-029-fascin (A) and Z1362873773-fascin (B) complexes. The graph shows the van der Waals contribution (VdW Energy, in teal color), electrostatic contribution (Elec. Energy, in plum color), and Total Energy (in amber color).

Finally, we determined the main non-bonded interactions obtained by MD simulations. At this point, most of them coincided with the docking calculation results. Two more residues were highlighted for the NP-G2-029 Fascin complex than the rest. Phe216 and Asp217 reveal high values of Total Energy, obtaining -29.1 and 23.5 kJ/mol for Van der Waals energy and -9.3 and -11.1 kJ/mol for electrostatic energy, respectively (Figure 14A). Regarding Z73 (Figure 14B), the interactions obtained followed a more even pattern. Several residues report total energy values between -10 and -20 kJ/mol. Among them, we can see some of them that match with the docking results and are known as crucial residues in the fascin union^29^, such as Phe14 (E_total= -13.3 kJ/mol2), Leu48 (E_total= -10.1 kJ/mol2), Trp101 (E_total= -17.9 kJ/mol2), Val134 (-11.9 kJ/mol2) and Phe216 (-15.7 kJ/mol2).

**Figure 14.**
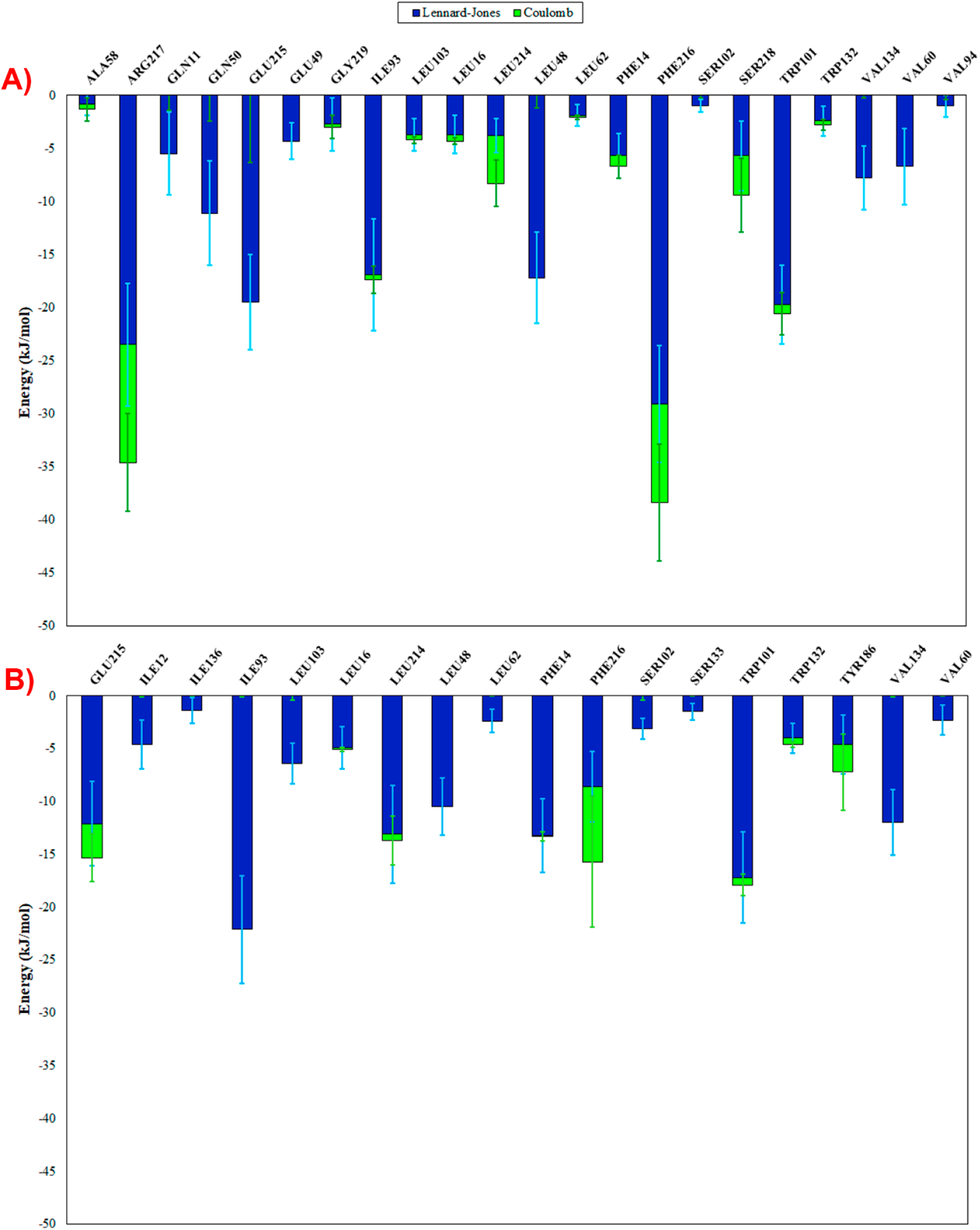
Non-bonded interaction analysis plots for NP-G2-029-fascin (A) and Z1362873773-fascin (B) complexes. The graph plots indicate the energy for each key residue split into different contributions (LJ (blue) and Coulomb (green)).

Consequently, the MD simulations correlated with the experimental results, showing Z1362873773 as a molecule binding to actin-binding site 1. The consensus of the RMSD, free binding energy validation by MMPBSA, and interaction analysis present a potential ligand with a stable and strong union with fascin, avoiding its function.

### Drug-like property predictions

By meticulously predicting the ADMET values for compound Z1362873773, we have provided a comprehensive understanding of its pharmacokinetic profile. This knowledge is crucial for its potential use in subsequent *in vivo* experiments and clinical trials. Our comparison with known fascin inhibitors, some of which have already been used as drugs, further solidifies the significance of our findings. Table 2 presents the predicted ADMET values for each molecule, setting the acceptable ranges that deem them favorable for a drug.

**Table 2.**
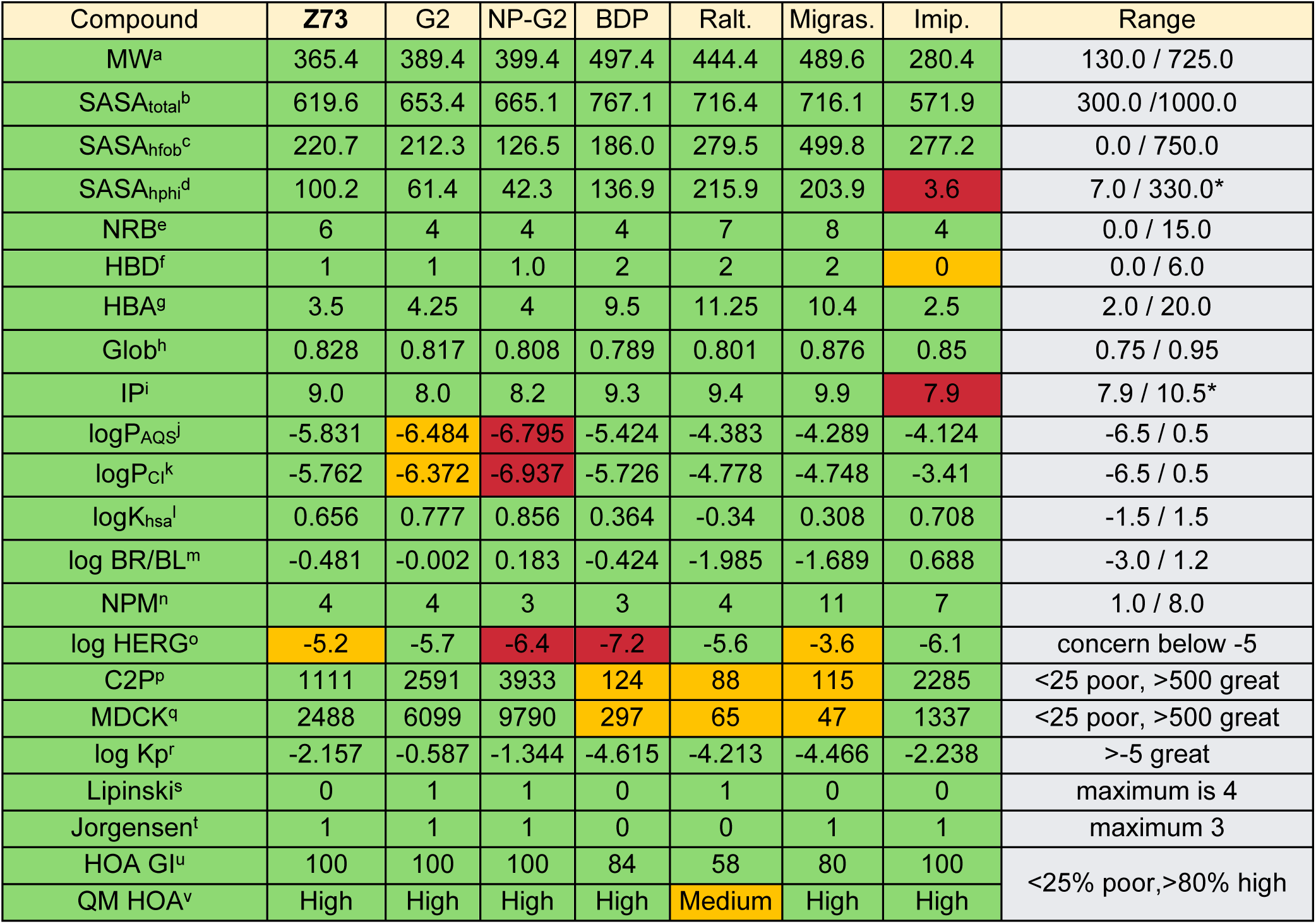
ADMET properties calculated by QikProp tools from MAESTRO for Z (Z73), G2, NP-G2-044 (NP-G2), BDP (BDP-13176), raltegravir (Ralt.), Migrastatin (Migras.), and imipramine (Imip). . Green indicates that the value is within the range, red indicates that is out of range. Orange indicates that the value is one of the limits of the range. aMolecular Weight (Da); bTotal solvent-accessible molecular surface (Å2); cHydrophobic solvent-accessible molecular surface (Å2); dHydrophilic solvent-accessible molecular surface (Å2); eNumber of rotatable bond; fHydrogen Bond Donated; gHydrogen Bond Accepted; hGlobularity (Sphere=1) iIonization Potential (eV) jlogP for aqueous solubility; klog P for conformation independent; llog of constant to human serum albumin; m log of predicted blood/brain barrier partition coefficient; nNumber of Primary Metabolites; olog IC50 value for HERG K+ Channel Blockage; pApparent Caco-2 Permeability (nm/sec); qApparent MDCK Permeability (nm/sec); rlog Kp for skin permeability; sNumber of violations for Lipinski rule; rNumber of violations for Jorgensen rule; uHuman Oral Absorption in gastrointestinal tract; vQualitative Model for Human Oral Absorption.

First, we discussed the predicted descriptors related to the bioavailability of Z1362873773. For this, we focused on the following properties of Table 2: the logP for aqueous solubility (logPAQS), the Apparent Caco-2 Permeability (C2P), the number of primary metabolites (NPM), the Jorgensen’s Rule of Three (Jorgensen) and the Lipinski’s Rules^35^. logPAQS is a critical value in this context because drug absorption in the body strongly depends on solubility. In this case, compound Z1362873773 showed a value of -5.831, which improved concerning the predictions for G2 and NP-G2-044 compounds. The number of primary metabolites (4) was also low enough not to affect bioavailability due to the production of metabolites, which could affect the bioavailability of the main compound. The compound violates only one Jorgensen rule and 0 for the Lipinski rule, which is a good result for its determination as a drug. Thus, in the four cases, the values predicted by QikProp for all four descriptors were within the ranges considered positive for a suitable pharmacokinetic profile, even improving those calculated for some already known fascin inhibitors.

Plasma protein binding is a critical factor for determining its effectiveness. It influences the concentration of free and active drugs in the blood and their distribution and removal from the body. Strong binding to plasma proteins such as human serum albumin can decrease drug availability for therapeutic purposes but can also prolong its effects by acting as a reservoir^36^. The log Khsa value provided important insights into these pharmacokinetic properties. Z1362873773 shows a value equal to 0.656. Thus, it falls within the established ranges to suggest that its interaction with plasma proteins is not sufficiently robust to impact the molecule’s availability and distribution^35^.

We also obtained favorable predicted values for human absorption. Qualitative Human Oral Absorption (HOA) and human oral absorption in the gastrointestinal tract (HOA GI) are notably high, reaching 100%. These findings suggest that Z1362873773 can be efficiently absorbed orally, indicating its potential as a drug candidate. Skin permeability is a potential route for the administration of several drugs. A crucial determinant in this regard is the QP log Kp for the skin permeability (log Kp) value, which indicates the ease of access of the molecule to the skin. For compound Z, QikProp predicted a QP log Kp value of -2.157. Typically, values approaching -2 suggest favorable permeability, indicating a probable entry route through the skin for this compound^37,38^. Finally, in this specific case, the drug should not cross the blood-brain barrier (BBB) because Z1362873773 is intended to act outside the central nervous system. QikProp calculates a predicted log BR/BL value, which indicates the ability of a compound to penetrate the BBB. The calculated value for the Z1362873773 compound is - 0.481. A negative log BR/BL value suggested a low likelihood of BBB penetration^35,39^.

Our predictions for ADMET properties by QikProp for Z1362873773 are within the expected range for a potentially effective drug and highlight its safety and efficacy. According to these predictions, Z1362873773 demonstrates favorable oral absorption and skin penetration. Its bioavailability is sufficient, and its plasma protein binding is weak enough not to interfere with its bioavailability significantly. Moreover, the predicted values suggest that Z1362873773 is unlikely to cross the BBB and cause adverse effects in the central nervous system. These findings, coupled with the fact that Z1362873773 displayed predicted descriptor values that were more promising than those of known inhibitors, instill confidence in its potential as a drug candidate. For example, significant improvements were observed in the solvent-accessible surface area (SASA), hydrogen bonding, and ionization potential, which are crucial for the interaction of the drug with the target and its surrounding environment^40^. Furthermore, Zitcompound showed favorable log HERG K+ channel values, suggesting a reduced risk of cardiac arrhythmias as a potential side effect^41^.

In our study on toxicity target prediction, we identified several potential targets for each of the two pharmacophoric models. Initially, we analyzed the hit molecules and their corresponding targets using LigandScout, based on a model incorporating all features (Table 3). Tasquinimod, a recognized antiangiogenic agent, has emerged as the primary hit. Although the literature references some immunomodulatory proteins, no specific target has yet been confirmed for Tasquinimod^42,43^. The other compounds identified using this method lacked notable targets associated with adverse effects. Overall, no targets associated with toxicity or adverse effects were detected in this model.

**Table 3.**
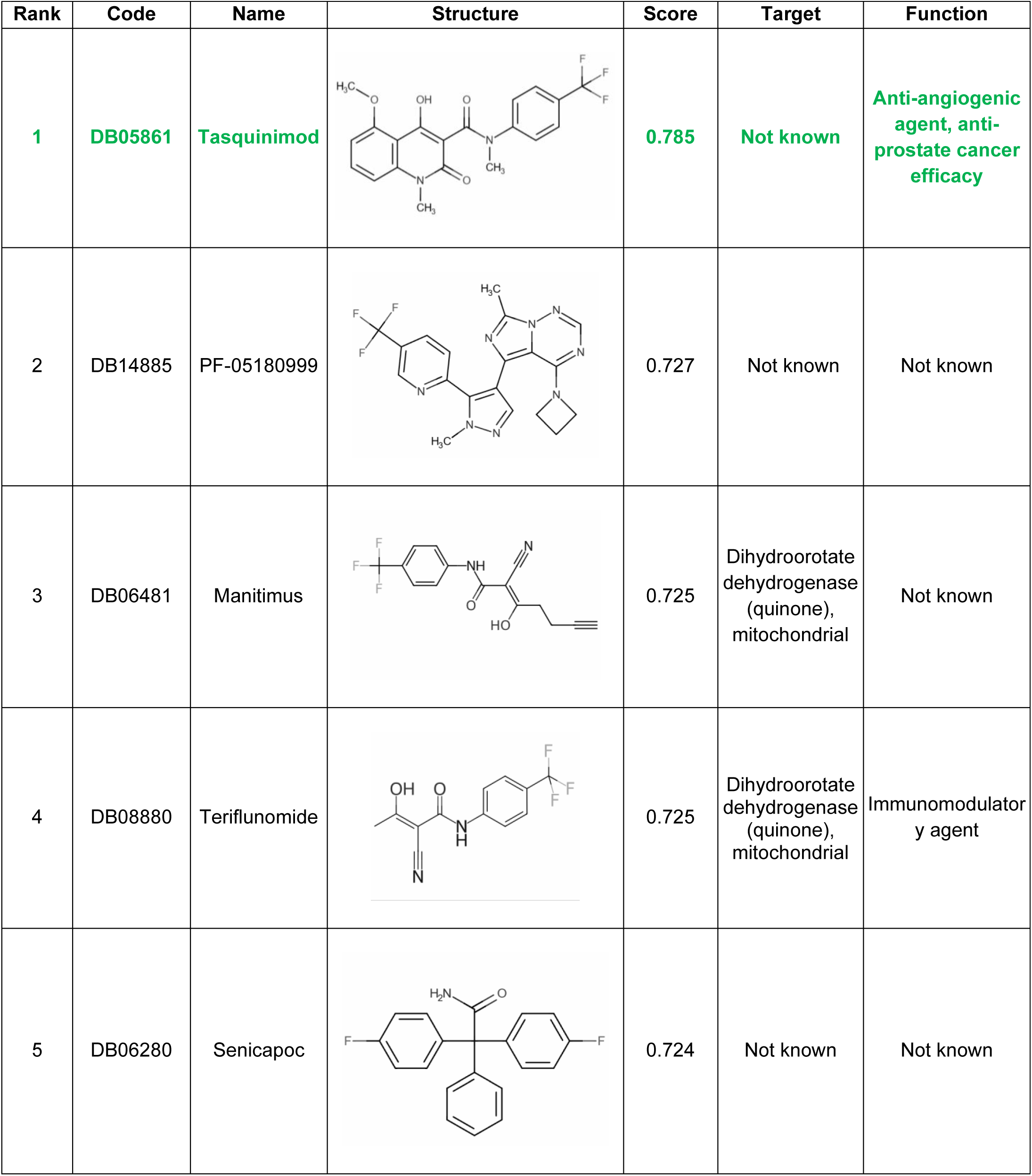
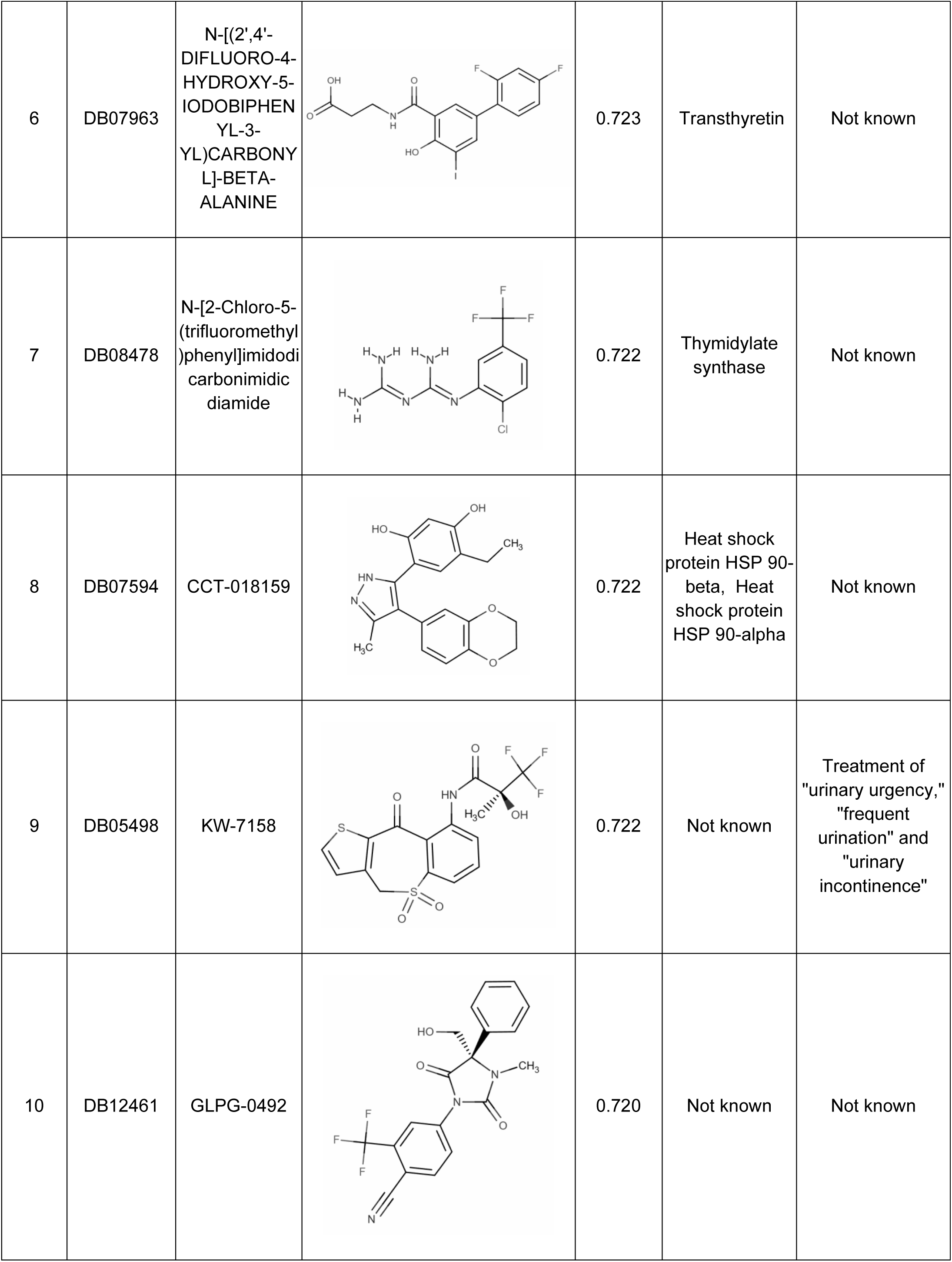
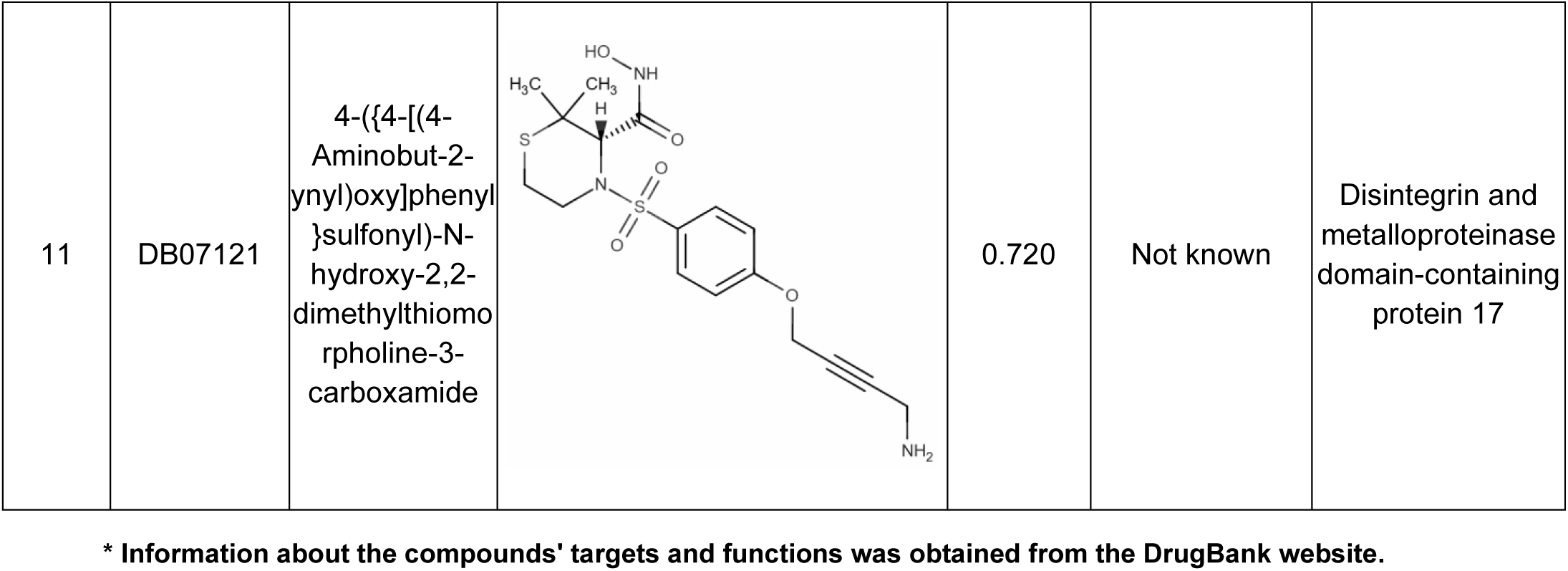
Results for Drug Target prediction obtained from the model generated with all the features found by Ligand Scout for the Z1362873733 compound.

| Rank | Code | Name | Structure | Score | Target | Function |
| --- | --- | --- | --- | --- | --- | --- |
| 1 | DB05861 | Tasquinimod |  | 0.785 | Not known | Anti-angiogenic agent, anti-prostate cancer efficacy |
| 2 | DB14885 | PF-05180999 |  | 0.727 | Not known | Not known |
| 3 | DB06481 | Manitimus |  | 0.725 | Dihydroorotate dehydrogenase (quinone), mitochondrial | Not known |
| 4 | DB08880 | Teriflunomide |  | 0.725 | Dihydroorotate dehydrogenase (quinone), mitochondrial | Immunomodulatory agent |
| 5 | DB06280 | Senicapoc |  | 0.724 | Not known | Not known |
| 6 | DB07963 | N-[(2',4'-DIFLUORO-4-HYDROXY-5-IODOBIPHENYL-3-YL)CARBONYL]-BETA-ALANINE |  | 0.723 | Transthyretin | Not known |
| 7 | DB08478 | N-[2-Chloro-5-(trifluoromethyl)<br>phenyl]imidodi<br>carbonimidic<br>diamide |  | 0.722 | Thymidylate<br>synthase | Not known |
| 8 | DB07594 | CCT-018159 |  | 0.722 | Heat shock<br>protein HSP 90-beta, Heat<br>shock protein<br>HSP 90-alpha | Not known |
| 9 | DB05498 | KW-7158 |  | 0.722 | Not known | Treatment of<br>"urinary urgency,"<br>"frequent<br>urination" and<br>"urinary<br>incontinence" |
| 10 | DB12461 | GLPG-0492 |  | 0.720 | Not known | Not known |
| 11 | DB07121 | 4-({4-[(4-Aminobut-2-ynyl)oxy]phenyl)sulfonyl)-N-hydroxy-2,2-dimethylthiomorpholine-3-carboxamide |  | 0.720 | Not known | Disintegrin and metalloproteinase domain-containing protein 17 |

\* Information about the compounds' targets and functions was obtained from the DrugBank website.

Our analysis using the Z1362873733 model, derived from the G2 compound model features (Table 4), NP-G2-044, a derivative of the Genin compound structure, stands out as the top candidate in our calculations. Furthermore, we have come across various other molecules that serve different functions, including acting as androgen receptor modulators, antidepressants, and even treatments for other types of cancer, such as Enasideb for acute myeloid leukemia (AML). Nonetheless, we have not encountered any targets or functions within these identified compounds that could be considered harmful or raised concerns regarding potential side effects.

**Table 4.**
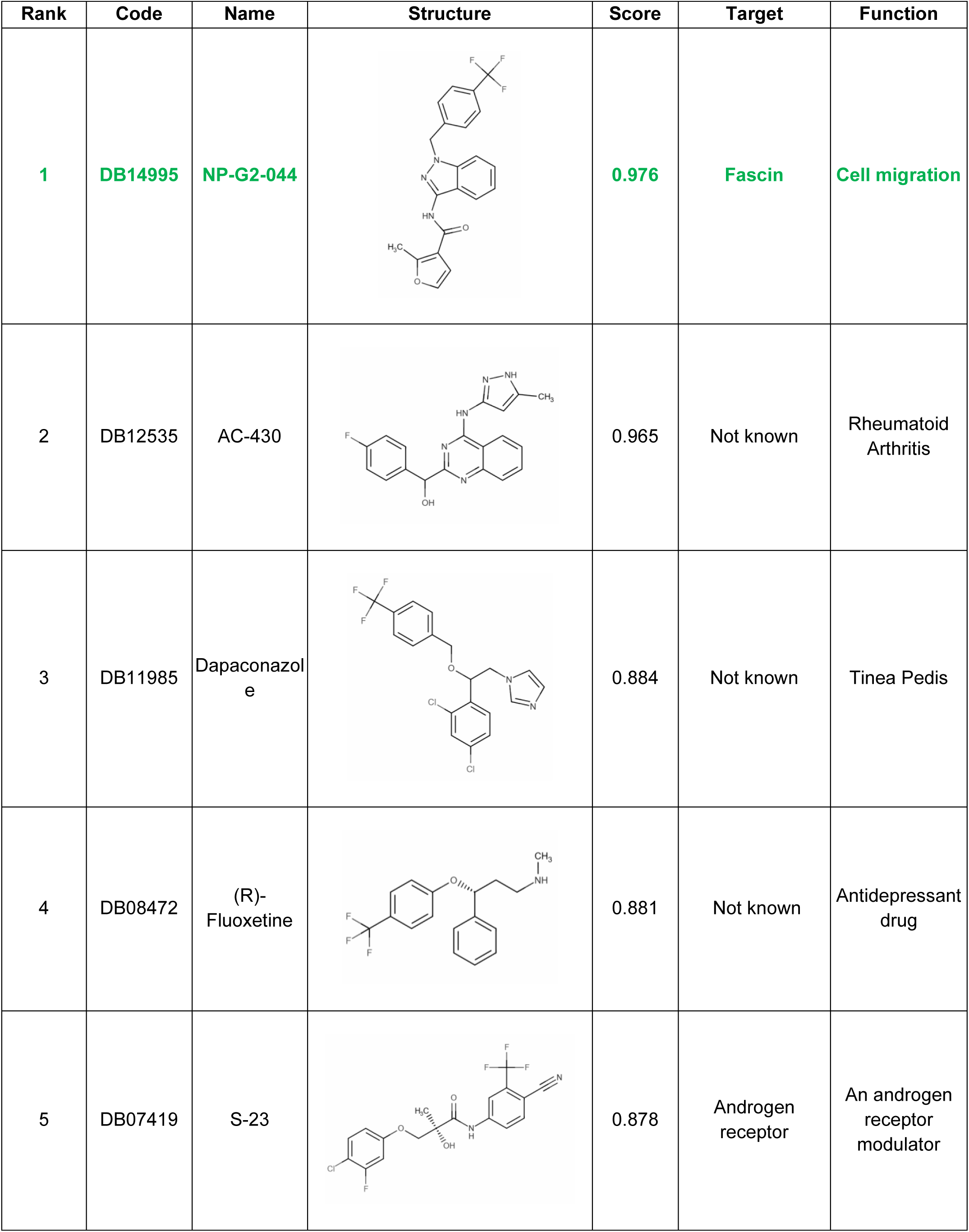
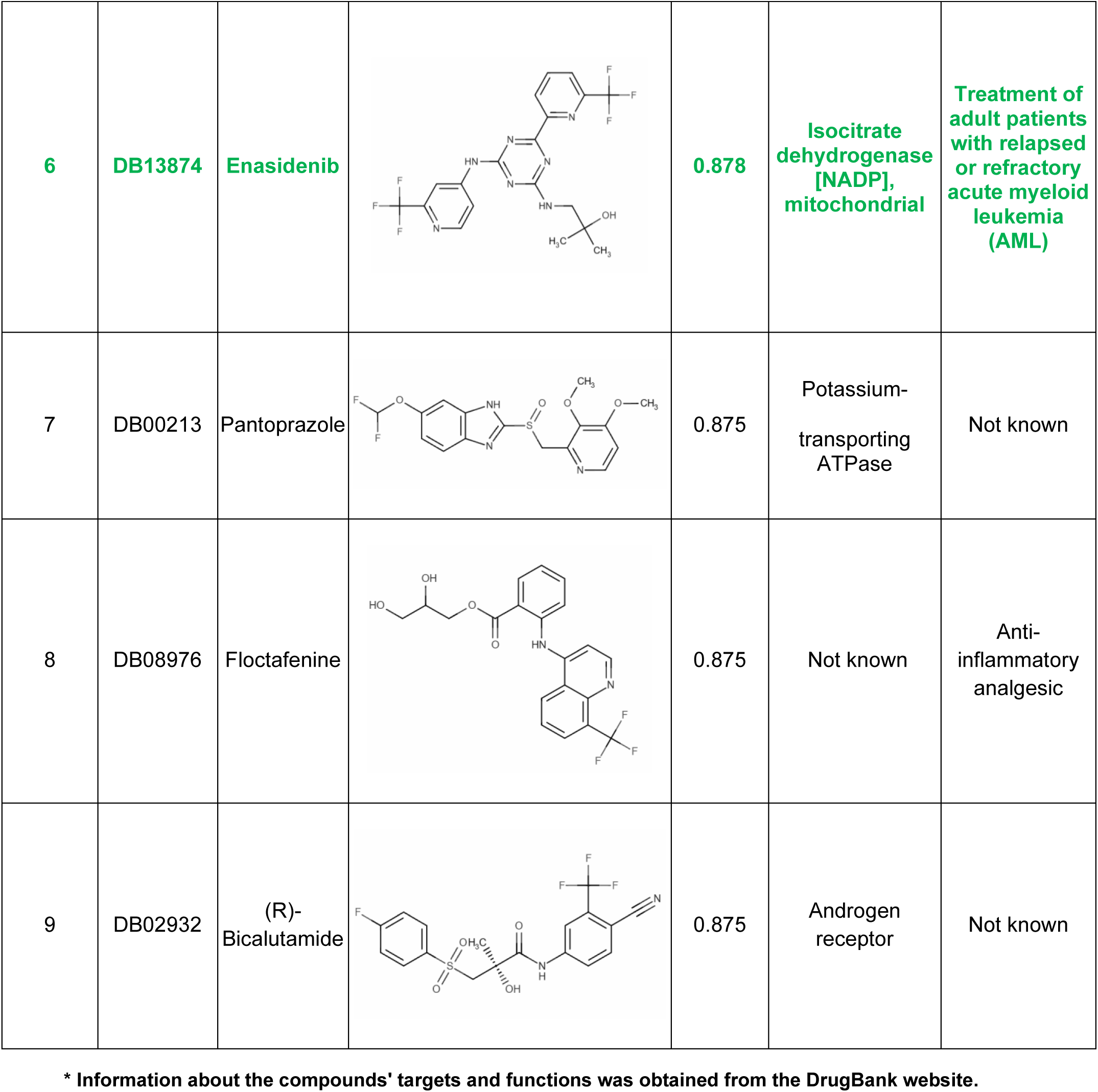
Results for Drug Target prediction obtained from the model generated just from the features that coincide with the G2 pharmacophore model.

| Rank | Code | Name | Structure | Score | Target | Function |
| --- | --- | --- | --- | --- | --- | --- |
| 1 | DB14995 | NP-G2-044 |  | 0.976 | Fascin | Cell migration |
| 2 | DB12535 | AC-430 |  | 0.965 | Not known | Rheumatoid Arthritis |
| 3 | DB11985 | Dapaconazole |  | 0.884 | Not known | Tinea Pedis |
| 4 | DB08472 | (R)-Fluoxetine |  | 0.881 | Not known | Antidepressant drug |
| 5 | DB07419 | S-23 |  | 0.878 | Androgen receptor | An androgen receptor modulator |
| 6 | DB13874 | Enasidenib |  | 0.878 | Isocitrate dehydrogenase [NADP], mitochondrial | Treatment of adult patients with relapsed or refractory acute myeloid leukemia (AML) |
| 7 | DB00213 | Pantoprazole |  | 0.875 | Potassium-transporting ATPase | Not known |
| 8 | DB08976 | Floctafenine |  | 0.875 | Not known | Anti-inflammatory analgesic |
| 9 | DB02932 | (R)-Bicalutamide |  | 0.875 | Androgen receptor | Not known |

\* Information about the compounds' targets and functions was obtained from the DrugBank website.

## Discussion

Fascin is one of the main target proteins involved in the migration and propagation of colorectal cancer. This protein is overexpressed in different types of cancer, especially in those with poor prognosis and more aggressive invasion. Several studies have elucidated possible molecules that can inhibit fascin function to date. Nevertheless, the chemical space explored in this context is still limited. This study explored a more extensive chemical space using the Enamine HTS library, which contains more than one million compounds, to identify potential inhibitors of fascin.

We completed a workflow to filter the entire library and search for potential compounds. The workflow starts with a ligand-based virtual screening, where we calculate the pharmacophoric similarity between G2, a known fascin inhibitor, and all the compounds from the library. Next, we validated the interaction of these compounds with fascin through biophysical in vitro assays. The next step was to determine the effect of the compound on migration and viability via assays in colorectal cancer cell lines and organoids. Finally, we showed evidence through in silico shreds of evidence obtained by docking and MD calculations of the complex formed between the final hit and fascin.

As demonstrated by the results of this case, our study’s protocol holds promise for identifying more potential compounds. Moreover, this workflow could be applied in various contexts, including those with new potential or even with larger libraries than the HTS Enamine library. This could significantly enhance the ability to identify compounds with high activity against specific backgrounds, potentially revolutionizing cancer treatment and drug development.

From our workflow, we isolated a novel compound, Z1362873773, which shares similarities with the known fascin inhibitor, G2. This synthetic molecule binds to fascin, inhibiting the formation of actin bundles crucial for cancer cell migration. In the migration and viability assays, Z1362873773 demonstrated values comparable to those of G2 in both *in vitro* and *ex vivo* models. Furthermore, the predicted ADME properties and drug targets suggest that Z1362873773 possesses ideal characteristics for drug action, marking a significant step forward in potential cancer treatments.

## Conclusions

In this study, we developed an efficient Ligand-Based Virtual Screening workflow, which led to the discovery of a novel anti-fascin agent from a large-scale combinatorial library (Enamine). Unlike previous studies, which used smaller libraries of FDA-approved compounds, our approach explored a more comprehensive chemical space. Starting with the pharmacophoric features of the G2 compound, we identified a new fascin inhibitor with a unique chemistry that is potentially free of intellectual property. Additionally, we provide molecular insights into how these compounds interact structurally with fascin, suggesting that this protocol could be applied to discover other anti-fascin agents.

## Patents

The authors declare that this work is included in a patent application (application number: EP24382860.5).

## Acknowledgments

This work was supported by a grant from the Scientific Foundation of the Spanish Association Against Cancer (Predoctoral Fellowship). This research was funded by JUNTA ANDALUCIA GRANT, the Andalusian Regional Government through the Grant: Proyectos de Excelencia (P18-RT-1193). The authors acknowledge the computing resources and technical support provided by the Plataforma Andaluza de Bioinformática at the University of Málaga. Powered@NLHPC research was partially supported by the super-computing infrastructure at the NLHPC (ECM-02).

## Abbreviations

ADMET: Absorption, distribution, metabolism, excretion, and toxicity
DSF: Differential Scanning Fluorimetry
DMSO: dimethyl sulfoxide
HTS: High-throughput Screening
MD: Molecular Dynamics
MMPBSA: Molecular Mechanics/Poisson Boltzmann Surface Area
SASA: Solvent accessible molecular surface
Tm: melting temperature

## Institutional Review Board Statement

The study was conducted in accordance with the Declaration of Helsinki, and approved by the Ethics Committee of the Santa Lucía University General Hospital (protocol code: REVERT, date of approval: 06/02/2021).

